# Variance Analysis of LC-MS Experimental Factors and Their Impact on Machine Learning

**DOI:** 10.1101/2023.05.01.538996

**Authors:** Tobias Greisager Rehfeldt, Konrad Krawczyk, Simon Gregersen Echers, Paolo Marcatili, Pawel Palczynski, Richard Röttger, Veit Schwämmle

## Abstract

**Background:** Machine learning (ML) technologies, especially deep learning (DL), have gained increasing attention in predictive mass spectrometry (MS) for enhancing the data processing pipeline from raw data analysis to end-user predictions and re-scoring. ML models need large-scale datasets for training and re-purposing, which can be obtained from a range of public data repositories. However, applying ML to public MS datasets on larger scales is challenging, as they vary widely in terms of data acquisition methods, biological systems, and experimental designs.

**Results:** We aim to facilitate ML efforts in MS data by conducting a systematic analysis of the potential sources of variance in public MS repositories. We also examine how these factors affect ML performance and perform a comprehensive transfer learning to evaluate the benefits of current best practice methods in the field for transfer learning.

**Conclusions:** Our findings show significantly higher levels of homogeneity within a project than between projects, which indicates that it’s important to construct datasets most closely resembling future test cases, as transferability is severely limited for unseen datasets. We also found that transfer learning, although it did increase model performance, did not increase model performance compared to a non-pre-trained model.

## Background

Large-scale studies of proteomes are essential to our understanding of the biological processes within an organism. The leading technology for characterizing thousands of proteins is Liquid-Chromatography Mass Spectrometry (LC-MS), which enables high-throughput quantification of protein abundances in a biological sample [1,2] (Figure 1).

**Figure 1.**
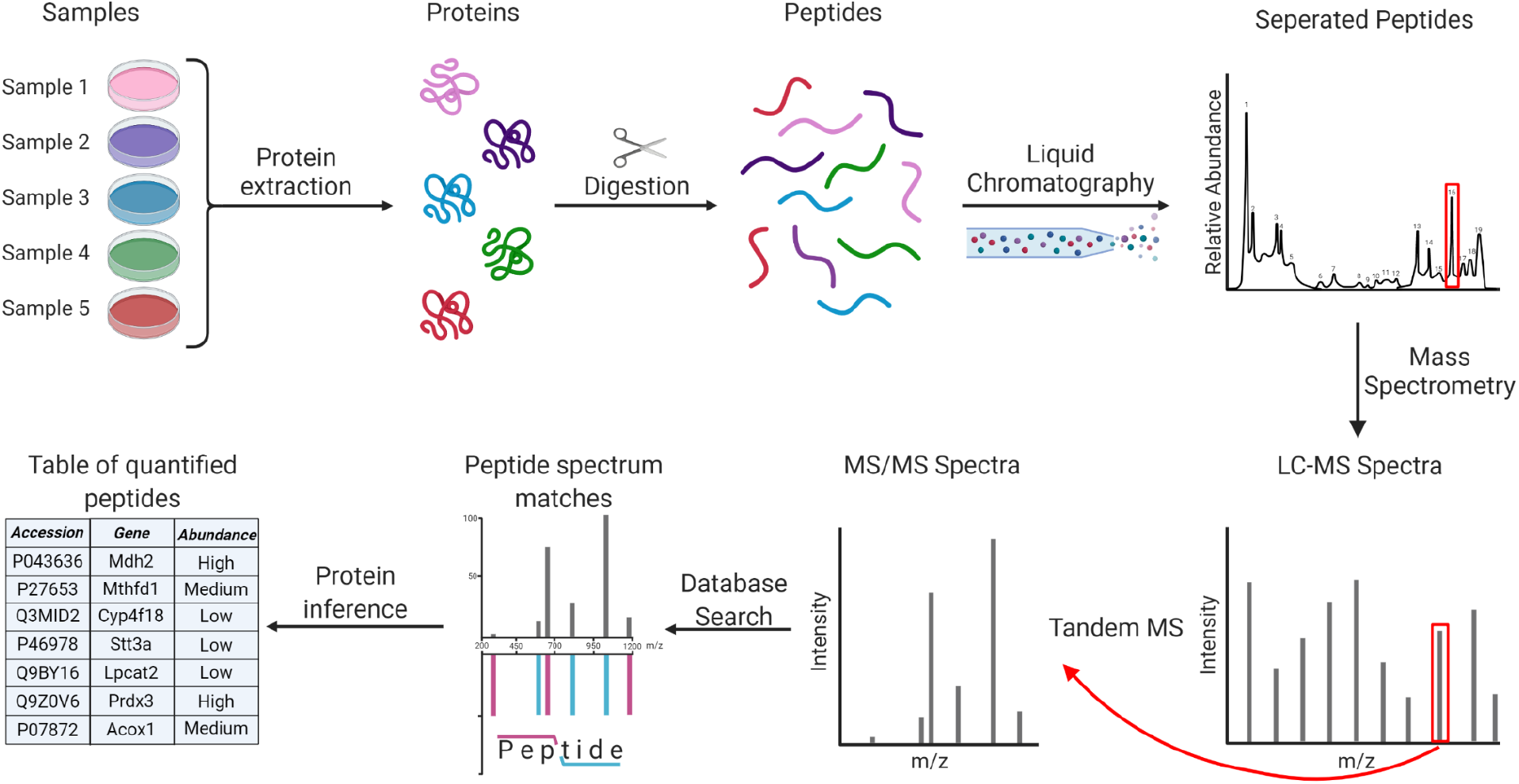
Simplified workflow of a Mass Spectrometry-based proteomics experiment. First, proteins are extracted from the biological samples, after which they are digested into peptides using enzymes, most often trypsin. Next, peptides are chromatographically separated and injected into the mass spectrometer where they are measured according to the mass over charge (m/z) and abundance (MS1). MS1 spectra from all peptide precursor ions are reported, and certain peptides are chosen for tandem-mass spectrometry (MS2), where they are fragmented along their amino acid backbone and identified by having their MS2 spectrum matched to a database of theoretical spectra. Lastly, peptides are quantified and summarized into proteins.[3–6]

LC-MS has become the standard within proteomics procedures and continues to generate vast amounts of data which, due to increasing demands from journals and reviewers, is often made publicly available in data repositories. This change has led to numerous public data sets being registered in online repositories such as the ProteomeXchange (PX) consortium [6]. The PXC contains references to over 17.000 projects, and its largest member, PRIDE, has more than a million raw files. Each raw file contains an average of 6.778 MS1 and 32.016 MS2 spectra, which amounts to over 39 billion mass spectra. These data repositories provide an invaluable resource for data repurposing to address novel biological questions or to benchmark new computational techniques for proteomics data analysis.

While efforts in harmonizing data accessibility within ProteomeXchange and standardizing the computational pipelines are ongoing [7], repurposing data from these repositories comes with a significant entry barrier, as they do not yet have any systematic criteria for metadata or data types.

Due to the advancements in machine learning (ML) model development, there is now an increasing interest in repurposing this rich LC-MS data to train complex ML models that can produce new insights and results not achievable by previous computational methods [8]. However, the large diversity of experimental procedures and biological systems requires careful consideration when applying bioinformatics methods to larger extracts of publicly available data as ML relies on careful balancing to reach optimal and correct performance.

Multiple ML algorithms and methods have been applied to MS data, such as regression models [9], random forest [10], and more recently neural networks [11]. Machine learning applications in proteomics are primarily focused on two aspects; (1) improving current methodologies such as database searches or de-novo sequences, or (2) predicting physico-chemical peptide properties such as LC-MS/MS spectra, retention time, or post-translational modifications (PTMs) [12–14]. Deep learning (DL) approaches function by generalizing the data, thereby generating distributions of the training data. However, due to the high complexity and large noise found in LC-MS data, many of the current approaches suffer from limited transferability, as they utilize synthetic, limited, or heavily stratified datasets for the purpose of training and testing their models [12,15,16]. These issues are further exacerbated by technical advances in the field such as ion mobility [17], which further increases the complexity and diversity of the data. The majority of current ML methods within computational proteomics also rely on unique and complex post-processing pipelines, such as peptide-specific indexed retention times (iRT) calculations, rendering the methods difficult to replicate and reducing their application range outside the original publication. One of the biggest shortcomings in machine learning, and particularly deep learning, is the problem of under and overfitting. These refer to situations in which a model either performs too well on the training data (overfitting) and generalizes poorly on unseen test data, or not well enough on the training data (underfitting) and subsequently also on unseen test data. Despite multiple attempts and the breadth of available data, these problems are still present in the field of predictive proteomics.

In this manuscript, we investigate the general reusability of public mass spectrometry data for machine learning applications, with a specific focus on potential pitfalls that could result in poor translatability to independently sampled data sets. We will do so by performing statistical analyses on the effect of the experimental setups on the variance of the generated data, and see how these effects impact the predictive capabilities of state-of-the-art deep learning models. This work is expected to have an impact on the data selection process in predictive proteomics, elevating the capabilities of current and future models, as well as highlighting the necessity for appropriate pre-processing and algorithmic choices.

## Data Description

For a comprehensive representation of publicly available MS data, we analyzed data from ∼60.500 raw files across ∼820 PRIDE projects, totaling ∼60 TB of raw files and metadata. 546 projects containing 33.426 raw files were used for neural network testing. All selected data had been previously analyzed with MaxQuant [18].

The full dataset was gathered from randomly sampled projects on PRIDE using MS2AI [19]. We restricted the retrieval to data from standard bottom-up proteomics experiments. We also did not have any initial queries on experimental or sample preparation, resulting in data from a wide breadth of sources from which we have sub-sampled smaller datasets for in-depth analyses (Table 1). In total, we gathered spectra and metadata for ∼151M individual unmodified peptides (Figure 2).

**Table 1.** Overview of datasets used in model training or testing.

| Name | Section | Minimal score | Peptide counts | Description |
| --- | --- | --- | --- | --- |
| *PT17* | 1 | 100 | 750.000 | Dataset constructed from PXD004732 |
| *PT19* | 1 | 100 | 750.000 | Dataset constructed from PXD010595 |
| *Limit* | 1 | 100 | 750.000 | 750.000 peptides excluding PXD004732 and PXD010595 |
| *Wide* | 1 | 150 | 2.000.000 | 2.000.000 peptides excluding PXD004732 and PXD010595 |
| *Long* | 2 | 150 | 2.000.000 | Gradient length equal to or above 100 minutes |
| *Short* | 2 | 150 | 2.000.000 | Gradient length equal to or below 60 minutes |
| *Lower* | 3 | 100 | 125.000 | Peptides with m/z values below 360 |
| *Upper* | 3 | 100 | 125.000 | Peptides with m/z values above 1300 |
| *Orbitrap* | Supp. | 150 | 2.000.000 | Instrument “orbitrap” according to PRIDE metadata |
| *Q. Exactive* | Supp. | 150 | 2.000.000 | Instrument “exactive” according to PRIDE metadata |
| *Human* | Supp. | 150 | 2.000.000 | “Human (Homo Sapiens)” from PRIDE metadata |
| *Mouse* | Supp. | 150 | 2.000.000 | “Mus musculus (mouse)” in PRIDE metadata |

All subset datasets are randomly sampled from the full ~151M peptide dataset, with Andromeda score and subset filters described in the “Minimal score” and “Description” columns respectively, no other filters were added when sampling. We also annotated the size of the subset dataset in the “Peptide count” column, with a 2M peptide count limit for each dataset.

**Figure 2.**
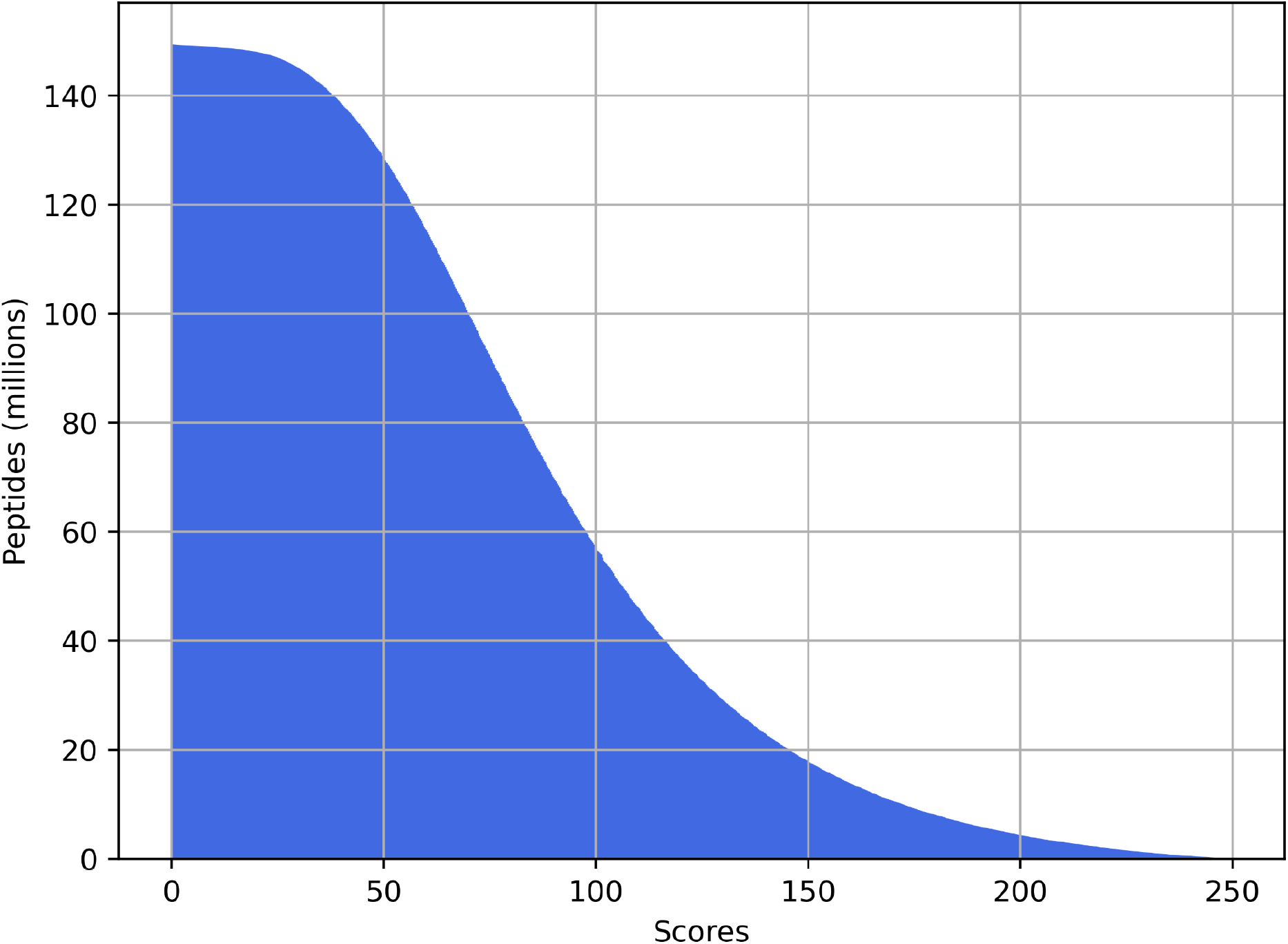
Andromeda score (MaxQuant) distribution plot of the ∼151M unmodified peptides in the complete data.

## Results and discussions

We performed a thorough statistical assessment of the data and trained multiple neural networks, in order to gauge the variance and evaluate the transferable capabilities of state-of-the-art models. However, in the field of predictive proteomics, and especially in retention time prediction, it is common to apply transfer learning to pre-trained models. This is done as a result of the poor transferability of the original networks, as the models do not achieve setup independence by being constrained to the experimental settings of the training data. However, transfer learning requires a large amount of data in a format suitable for machine learning, as well as significant computational expense, making it both data- and computationally-intensive. Additionally, it also requires expertise in both machine learning and programming. In order to test the effectiveness of transfer learning in predictive proteomics, we exhaustively transferred all of the trained models to every other dataset in the same section to assess the impact of transfer learning on performance.

All model metrics are measured in RTΔ, which measures the average time difference between predicted RT values and actual values.

### 1. Single vs multi-project variance

In our model comparisons, we found that the models trained on the *PT17* and *PT19* datasets performed significantly better than the models trained on the *Limit* and *Wide* datasets during both training and validation (Figure 3). Interestingly, the *PT* models also outperformed the *Wide* model, and performed comparably to the *Limit* model, when testing on the wide and limited datasets.

**Figure 3.**
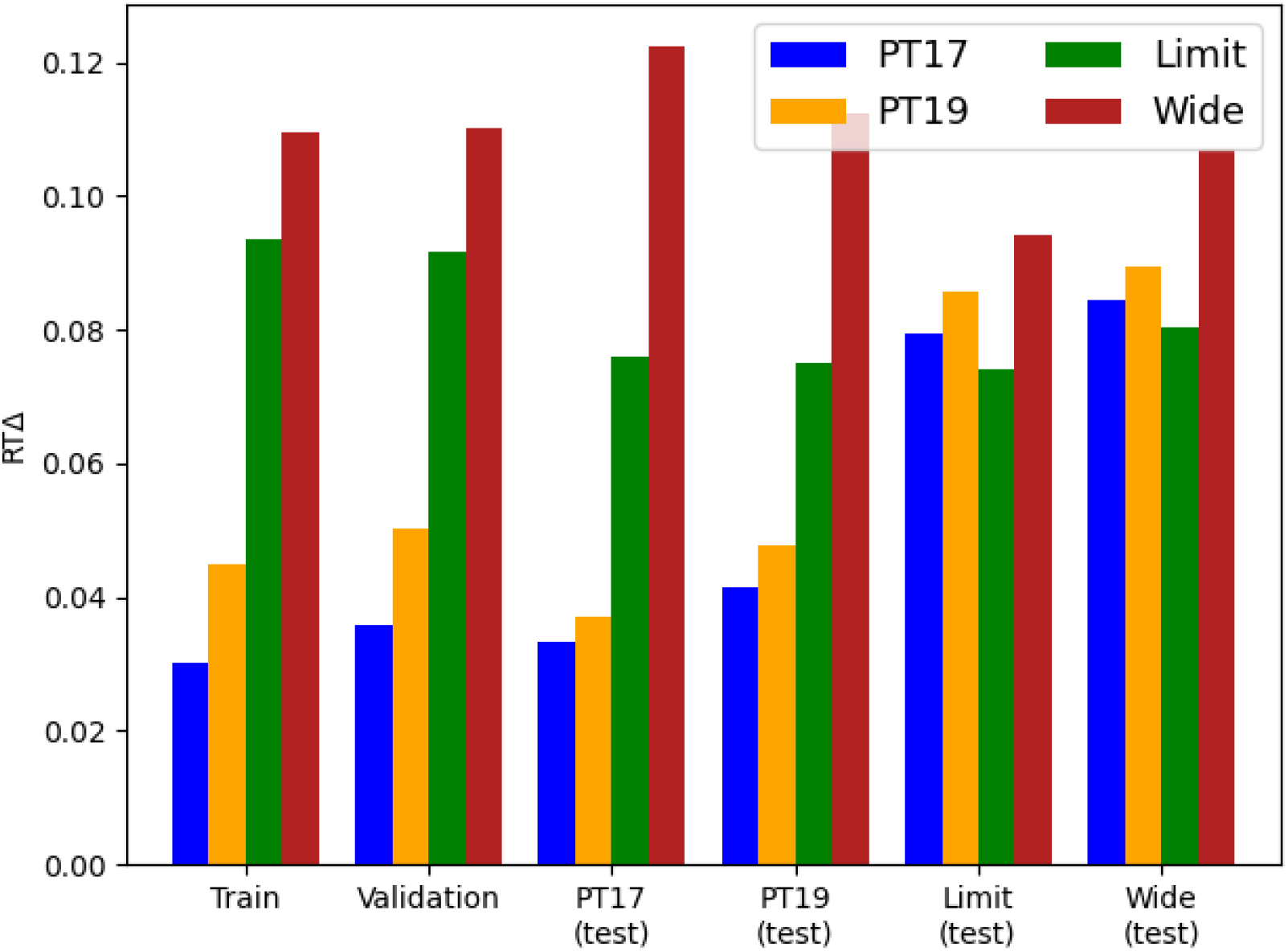
General variance model performance comparison. Each model was trained and validated on its original source datasets and then cross-tested for all test datasets. We compared the training and validation for all models, as well as cross-testing datasets and their respective model performance in terms of RTΔ

While we expected the training and validation of the *PT* models to outperform the *Wide* and *Limit* model, we did not expect the *PT* models to outperform the *Wide* and *Limit* models on the *Wide* and *Limit* test datasets. Furthermore, we also found that the *Limit* model outperformed the *Wide* model for all test cases, suggesting that increasing the amount of data and using stricter scoring criteria does not necessarily improve the performance of trained models, and may even cause the models to underfit. One possible explanation for this is that longer and more complex peptides, which are easier to detect and in general receive higher peptide identification scores, also exhibit more variance in their elution times. We tested this and found that the peptides in both *PT* datasets have average peptide lengths of ∼12, with the *Limit* datasets containing a longer average peptide length of ∼13, and the *Wide* dataset containing an even longer average length of ∼14.

It is also possible that the reduced performance observed in the randomly sampled datasets is due to the presence of multiple variance-inducing factors in the experimental setup. These factors, such as the type of MS instrument or the selected species, are often difficult or impossible to account for when relying on large bulks of public data. Changing them can result in considerably different model performances (Supplementary Figure 1-2). In contrast, the *PT* datasets were measured under the same conditions on synthetic peptides, which reduces the presence of such variance-inducing factors. Additionally, the *PT* datasets have the advantage of using one single gradient length, while the limited and wide datasets use multiple different gradient lengths.

In order to evaluate their transferability, we applied transfer learning to all four models and found that, while some of the models improved performance compared to previous external testing, they mostly performed similarly to non-transferred models on the same dataset (Figure 4). In the case of the *Wide* dataset, transfer learning actually resulted in reduced performance for all transferred models. This suggests that the *Wide* dataset has significantly higher levels of heterogeneity between training and testing data compared to the other datasets. While these results show that transfer learning can be beneficial in certain scenarios, they also showed that most of the models simply improved or regressed to the transfer dataset. However, even if transfer learning did not provide significant predictive benefits, it did reduce the time needed to train the models by an average of 5.5% by converging faster (Supplementary Table 2).

**Figure 4.**
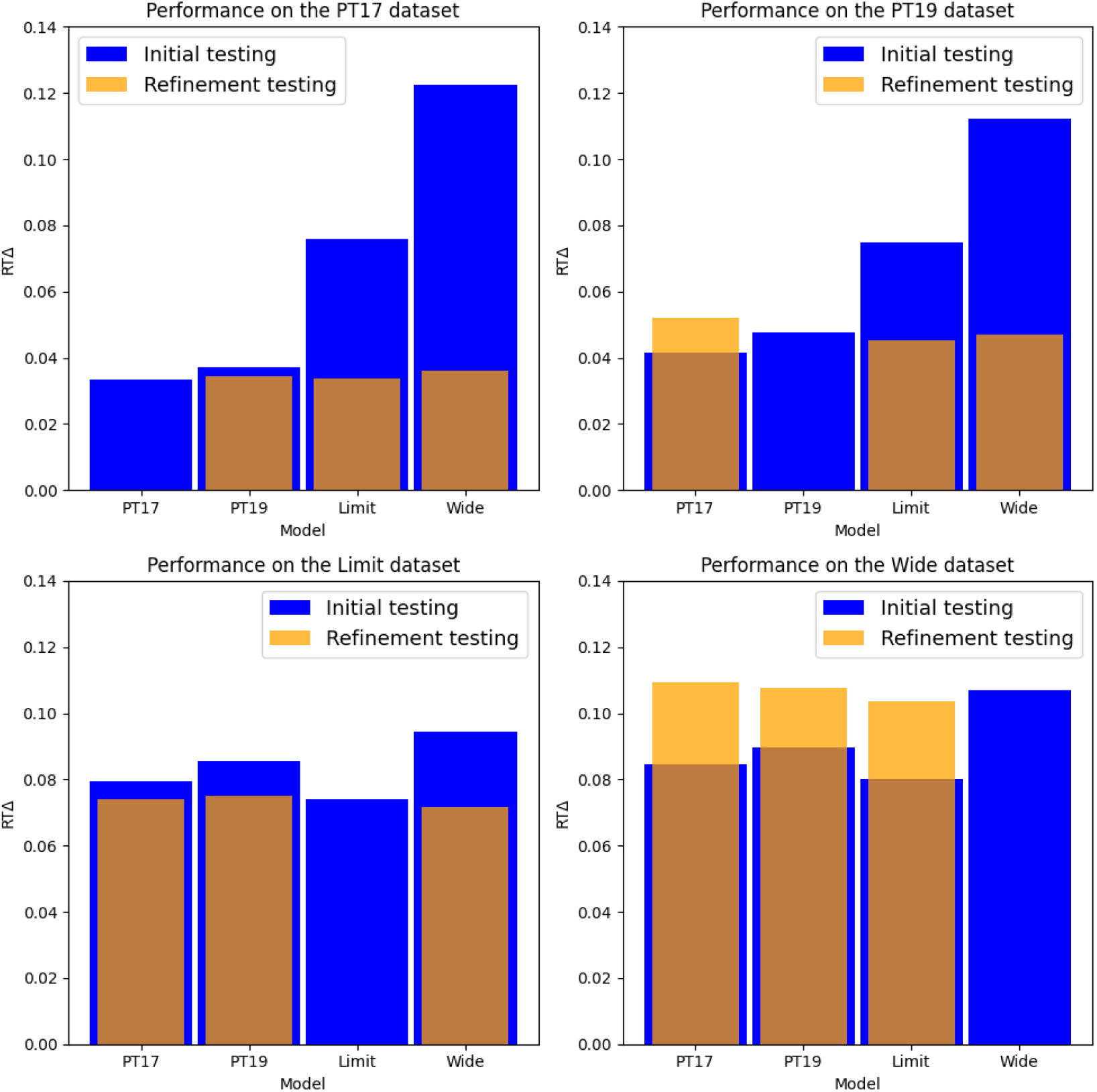
General variance transfer model performance comparison. Each model was transferred to each of the external datasets and re-tested on all four datasets. Datasets are separated by plots, denoting the performance difference of each model when trained on or transferred to identical datasets. Each bar has the original test metric in blue, and the transfer learning test metric overlaid in orange.

### 2 Gradient lengths

The retention time of a peptide in an LC-MS/MS system is determined by the interactions between the peptide and the stationary and mobile phases of the liquid chromatography system. In identical setups, the retention time of a peptide is considered reproducible[9].

Plotting the distribution of gradient length for all raw files with an overlaid cumulative distribution function (Figure 5a), we observe significant peaks at 60, 90, and 120 minutes, with ∼60% of all gradients being 0–120 minutes in length and the longest gradient being 800 minutes. While we did find single projects with as many as 15 different gradients, we also found that 70% of the 820 projects kept the same gradient length for all files, while only ∼5% employed more than two unique gradients (Figure 5b), indicating high levels of consistency in instrument configurations within individual projects.

**Figure 5.**
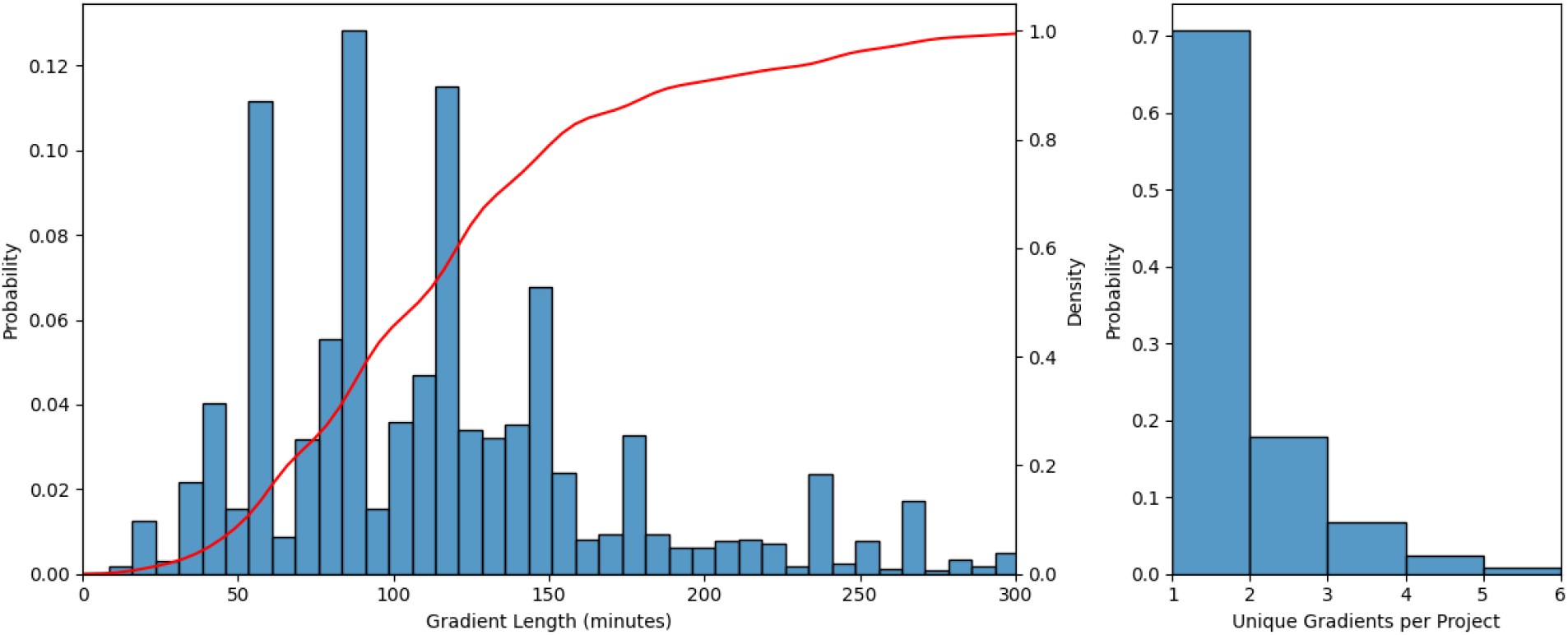
Gradient length distribution and unique gradient per project. We illustrated the gradient distribution of all 60.000+ raw files overlaid with the cumulative distribution (A). Also illustrated is the number of unique gradients found within any of the 820 projects (B)

The results of our deep learning use case showed that the *Short* gradient model performed significantly better than the *Long* gradient model (Supplementary Figure 3). This is further supported by its decreased performance on the *Long* gradient test dataset compared to the *Short* model.

These findings suggest that peptides from longer gradients generally express higher variance compared to peptides from shorter gradients, even after attempted peptide normalization. It also indicates that our normalization method for the effective gradient, which aims to mimic the linear iRT calculations used in the original Prosit paper, may not be effective for all gradients and raw files, reiterating the necessity of targeted post-processing pipelines.

Performing inference dropout on all models in the previous section shows that all of the models exhibit significantly higher uncertainty for the earliest and latest eluted peptides compared to those eluted closer to the middle of the gradient (Figure 6). Additionally, we observe that the *PT* models show a more linear prediction gradient than the *Limit* and *Wide* models, further suggesting the controlled nature of the ProteomeTools datasets outputs peptides in a more linear gradient, which fits better for our first-last peptide gradient normalization. We also observe less overall uncertainty in the PT models, likely due to the fact that they were trained on datasets with fewer peptides, lower average peptide retention times of approximately 32 minutes, and unified gradients, whereas the average retention times of the limit and wide datasets were significantly higher at 60+ minutes from multiple gradient lengths. This suggests that longer gradients lead to an increase in data variance and model uncertainty.

**Figure 6.**
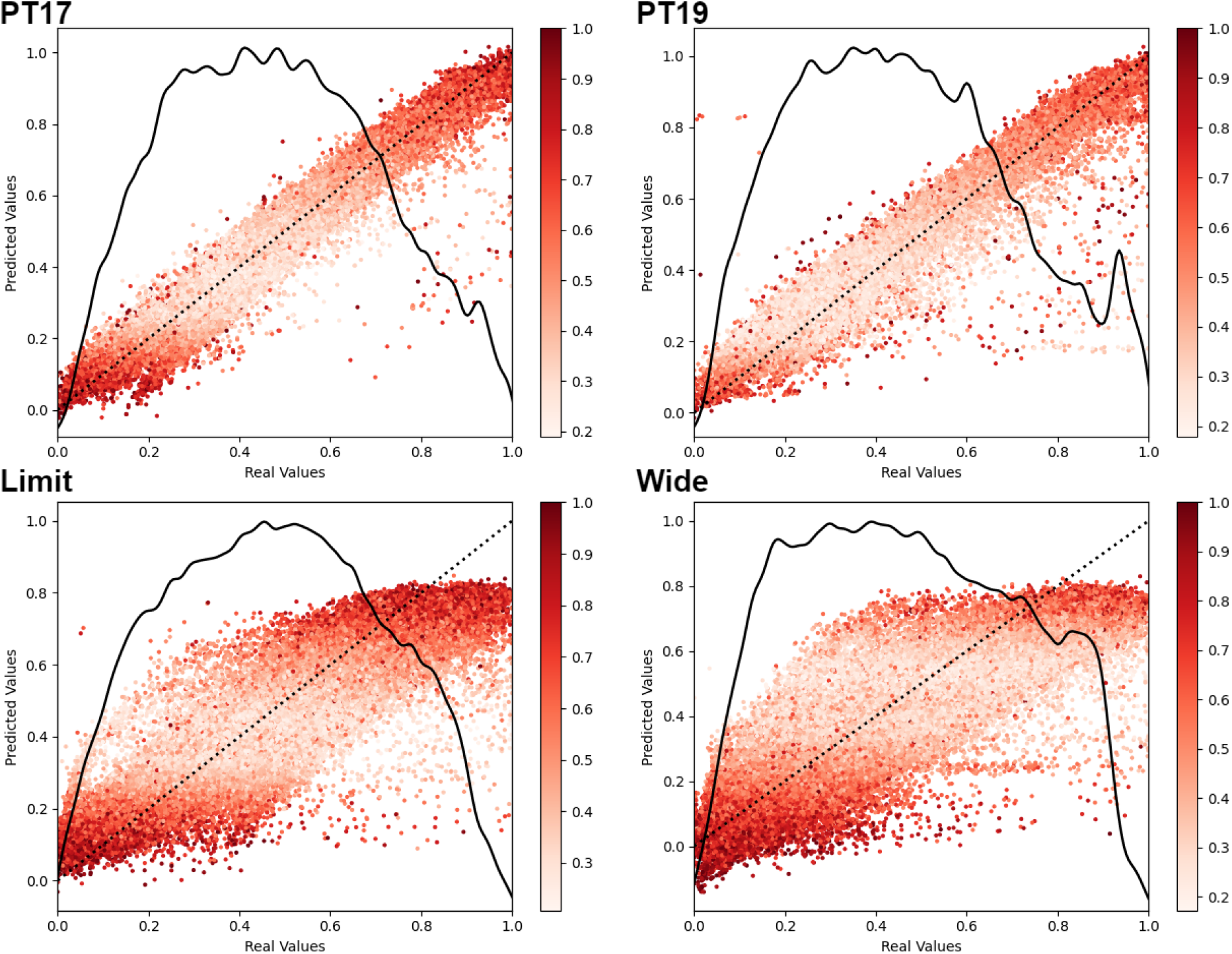
Bayesian model uncertainty estimates of general variance models. For each model; *PT17, PT19, Limit*, and *Wide*, we performed model inference on individual test datasets 25 times with dropout applied. The mean predicted retention times are plotted against the real retention time values with the normalized relative variance of each peptide illustrated in the colorization of the dots. The distribution of the underlying data points (line) are overlaid together with a y=x line (dot) for linearity comparison

Performing transfer learning on the gradient length models (Supplementary Figure 4) we observe that refining the *Long* model to the *Short* dataset resulted in a significant performance improvement. However, refining the Short model to the Long dataset did not result in any significant change in performance, although it still outperformed its non-transferred counterpart. Unlike what we observed in the previous section, transfer learning of the gradient length models came at an increase or stagnation in performance, indicating that the models retained information from the initial training datasets. We also note that transferred models took on average 25% longer to train compared to non-transferred models (Supplementary Table 3).

### 3 Mass-to-charge range filter

MS instruments have a range of setup parameters that can tailor the experiment to the needs of researchers. The m/z range filter is one of those parameters, as it restricts peptide data acquisition to a given m/z range. However, for data re-purposing, this range can also lead to a biased dataset for machine learning, as the sample might have contained a large range of peptides not reported by the instrument. When plotting the m/z filters of the mass acquisition range (Figure 7), we observe significantly more variance in the upper bound compared to the lower bound, meaning that our upper bound is highly correlated to the length of the filter. All violin plots exhibit a peak at one specific value, 350 for the lower bound, and 1500 for the upper bound, with a corresponding peak at 1150 for the filter lengths. These peaks correspond to the most commonly used filter which accounts for 33.9% of all raw files.

**Figure 7.**
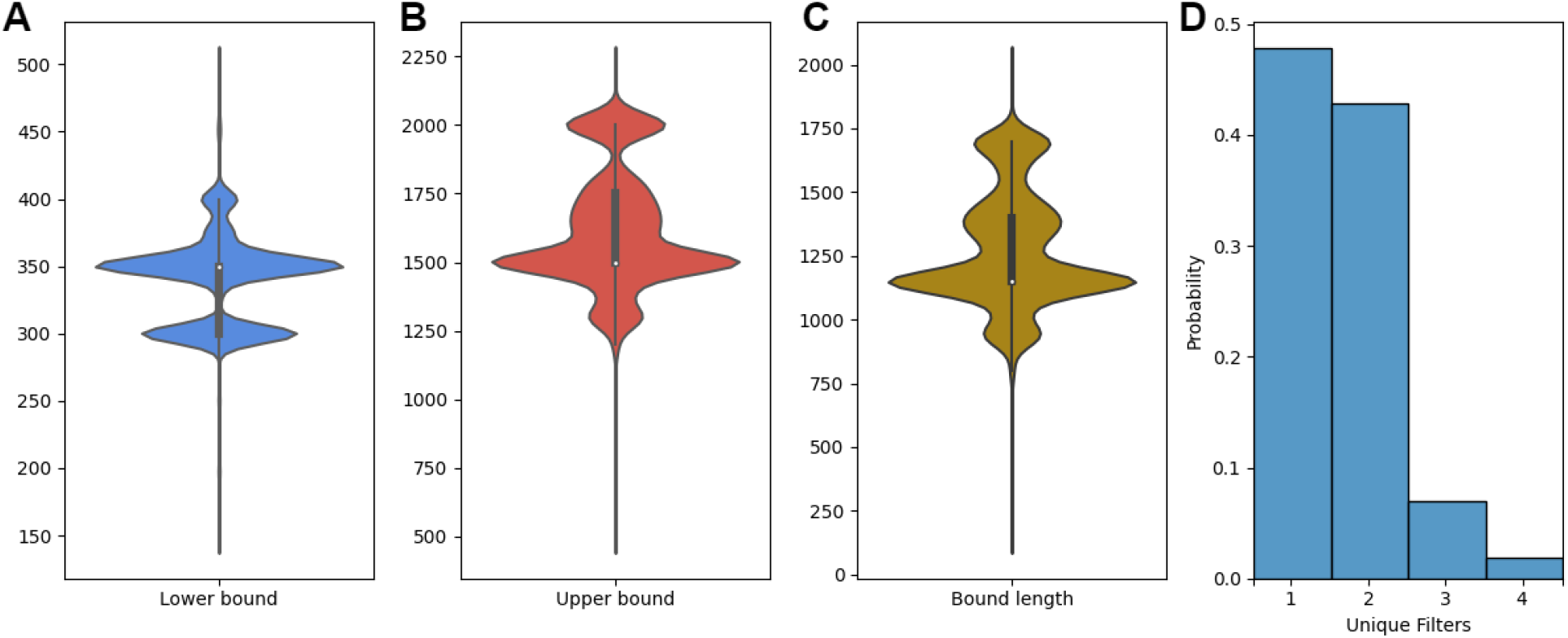
Illustration of the variability of m/z range filters found in different projects. Violin plots depicting the m/z filter bound distributions, with the lower bound plotted in blue (A), upper bounds plotted in red (B), and filter length plotted in yellow (C). A histogram of the total number of unique filters found within projects (D).

We also observed that 47% of all projects applied a single filter across all raw files, 43% of projects applied two filters and only 10% of the projects applied more than two unique filters. Consequently, there is significant homogeneity within a project while between projects the filters can differ considerably. If datasets are constructed using only one or a few unique filters, large portions of the data space may never be used for training, potentially limiting the transferability of the models (Supplementary Figure 5).

We also tested the model performance of the *PT17* and *PT19* models on peptide datasets only containing peptides outside of the original filter bounds (out-of-bounds, OOB, Supplementary Figure 6), and found that model performance was significantly worse when compared to the source test datasets. The OOB testing performance is also significantly worse when compared to *Limit* and *Wide* testing (*results and discussion section 1*), which also contained peptides in the OOB range. Interestingly, the models performed slightly worse on heavier OOB peptides than on lighter OOB peptides (Figure 8), despite lighter peptides exhibiting higher individual variance, and the distribution of lighter peptides being more concentrated at one end of the distribution compared to the heavier peptides (Supplementary Figure 7). Similarly to Figure 6, we again observe significantly more model uncertainty tied to earlier and later eluted peptides.

**Figure 8.**
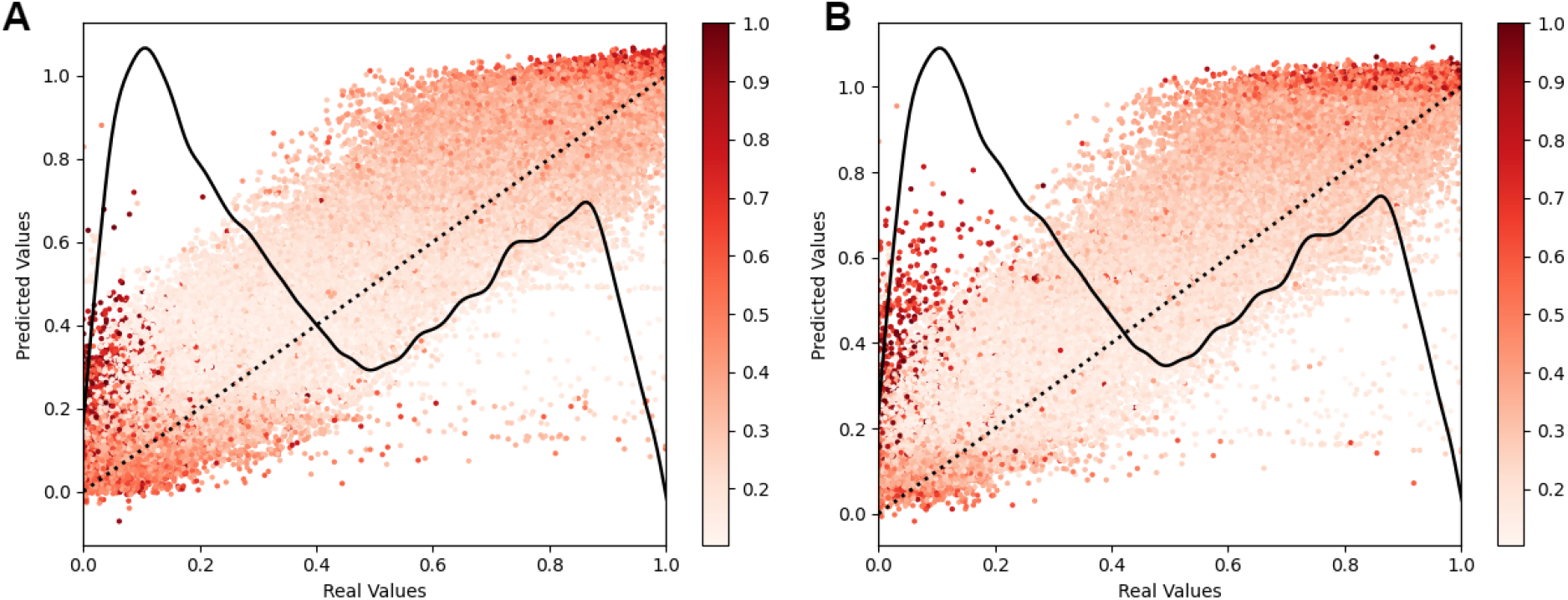
Bayesian model uncertainty estimates for out-of-bounds peptides. For *PT17* (A), and *PT19* (B) we performed model inference 25 times on peptides from datasets outside of the initial training datasets m/z bounds, with random dropout applied. The mean predicted retention times are plotted against the real retention time values with the normalized variance of each peptide illustrated in the colorization of the dots. The distribution of the underlying data points (line) are overlaid together with a y=x line (dot) for linearity comparison

The significant reduction in performance we observed in the OOB testing suggests that the models do not learn the underlying nature of AA weights, folding, and physico-chemical properties, factors which impact RT as well as the ionization and detectability [16] of peptides, as much as they memorize the retention times of certain peptide sequence patterns.

### 4 Fragmentation patterns

A perfect peptide fragmentation spectrum is a theoretical concept that consists of a discrete set of all characteristic peaks defined only by the peptide sequence. In reality, fragmentation spectra only contain subsets of these theoretical peaks with patterns based on the background contaminants from the instrument, fragmentation technique, collision energy, and more. In order to understand the challenges and limitations of the MS2 spectra for machine learning algorithms, we analyzed MS2 spectra peaks from 86 randomly sampled raw files containing more than 768 million peaks, as well as from three representative samples from the ProteomeTools 2019 dataset. This allows us to visualize not just the overall distributions found in MS2 spectra, but also how these distributions change and are affected by fragmentation techniques and activation energy. While the different techniques differ in activation energies; normalized collision energy (NCE) for CID and HCD, and charge state-dependent reaction time for ETD, we simplify both as the activation energy (AE) [20]. In hybrid fragmentation methods (ETciD and EThcD), the CID/HCD NCE is fixed while the ETD reaction time is varied.

When looking at the distribution of all m/z values found in the 86 randomly sampled files, we observe a clear bimodal distribution independent of peak selection or bin sizes; one distribution is located at ∼50-250 m/z, and the second distribution at ∼250-2000 m/z. The distribution at the lower m/z range disrupts the expected normally distributed peptide fragment ions found at 250-2000 m/z, and consists of clearly distinguishable high-density peaks corresponding almost exclusively to single amino acid residues (Fig. 9E and 9F) and some background noise. The singly charged amino acid residues are shown in the cumulative distribution plots in Figure 9, where amino acids are annotated if the m/z of the peak matches the collision ions *a, b*, or *y* or the electron-transfer ions *c* or *z*. The most abundant amino acid peaks are highlighted in Supplementary Table 1, and we observe that these peaks become more frequent at higher levels of peak selection (Supplementary Figures 8-9). It should be noted that the exact AA contribution to some peaks is uncertain, as multiple AA ions match the exact same m/z peaks.

**Figure 9.**
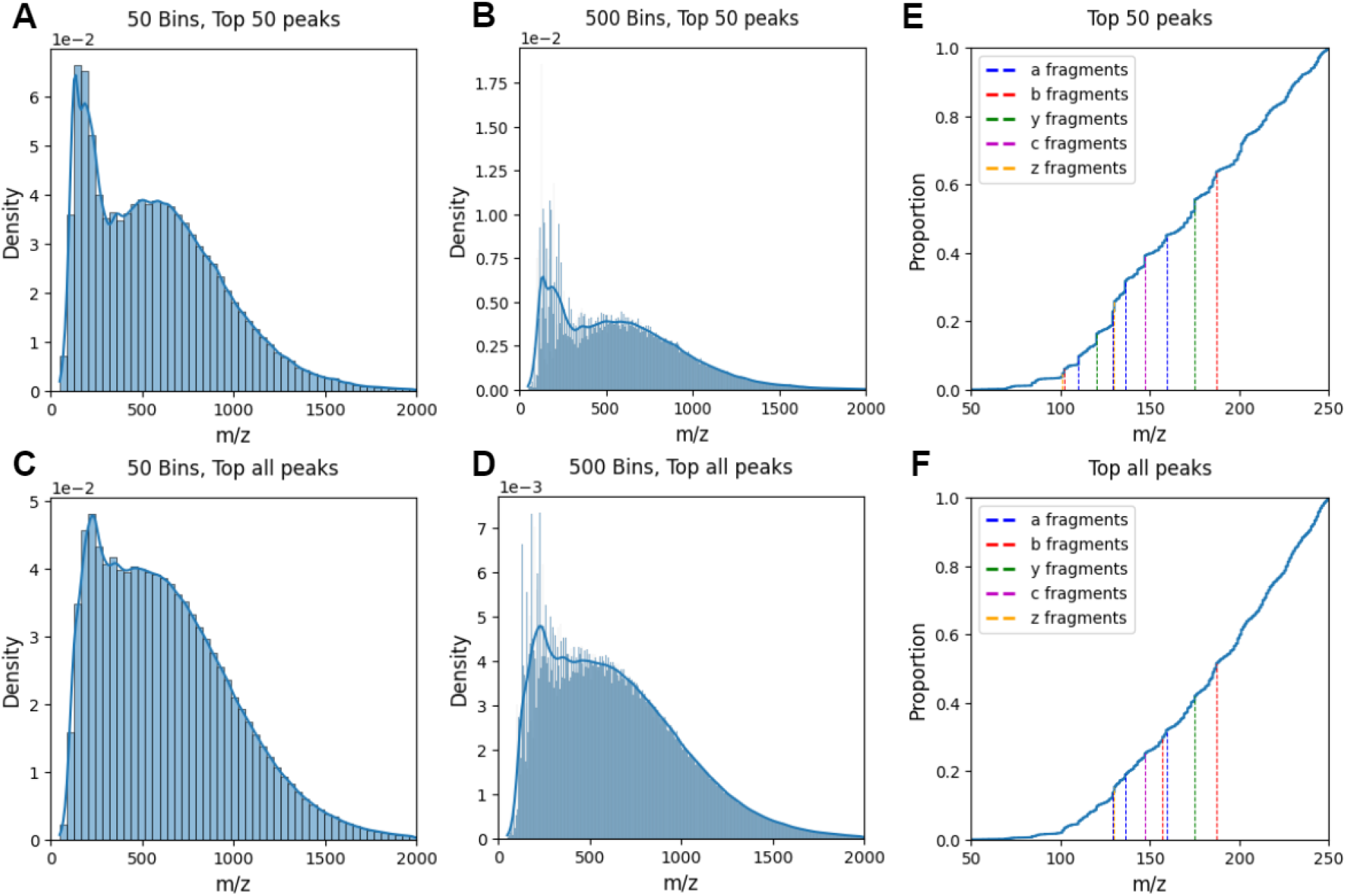
Illustration of how peak picking and peak binning affect the MS2 peak density plot and single amino acid density. The density distributions of peak m/z values shown were obtained without peak picking (C, D) and with the top 50 peaks (A, B). The spectra were then imposed into 50 (A, C) and 500 (B, D) bins. For each value of peak picking, we also illustrated the cumulative distribution plot of the peaks in 50-250 m/z with single AA residues overlaid. No stratification on the fragmentation method or its settings has been considered during this analysis.

When we compare the ProteomeTools files, we see that these single AA peaks are present in all fragmentation methods except CID (Figure 10, Supplementary Figures 10-11). Interestingly, the AA peak information gets weakened at different levels of activation energy for different files. In Supplementary Figures 12-13 these peaks are drastically reduced or almost non-existent in ETD variants at 32.27 ms reaction time, specifically, whereas Supplementary Figure 13 exhibits a more gradual loss of signal as activation energy is decreased. This suggests that more energy is needed to fragment sequences into single AAs, and without sufficient energy, these signals are weakened. The potential use of these ions will heavily depend on the applied fragmentation method and activation energies of the system.

**Figure 10.**
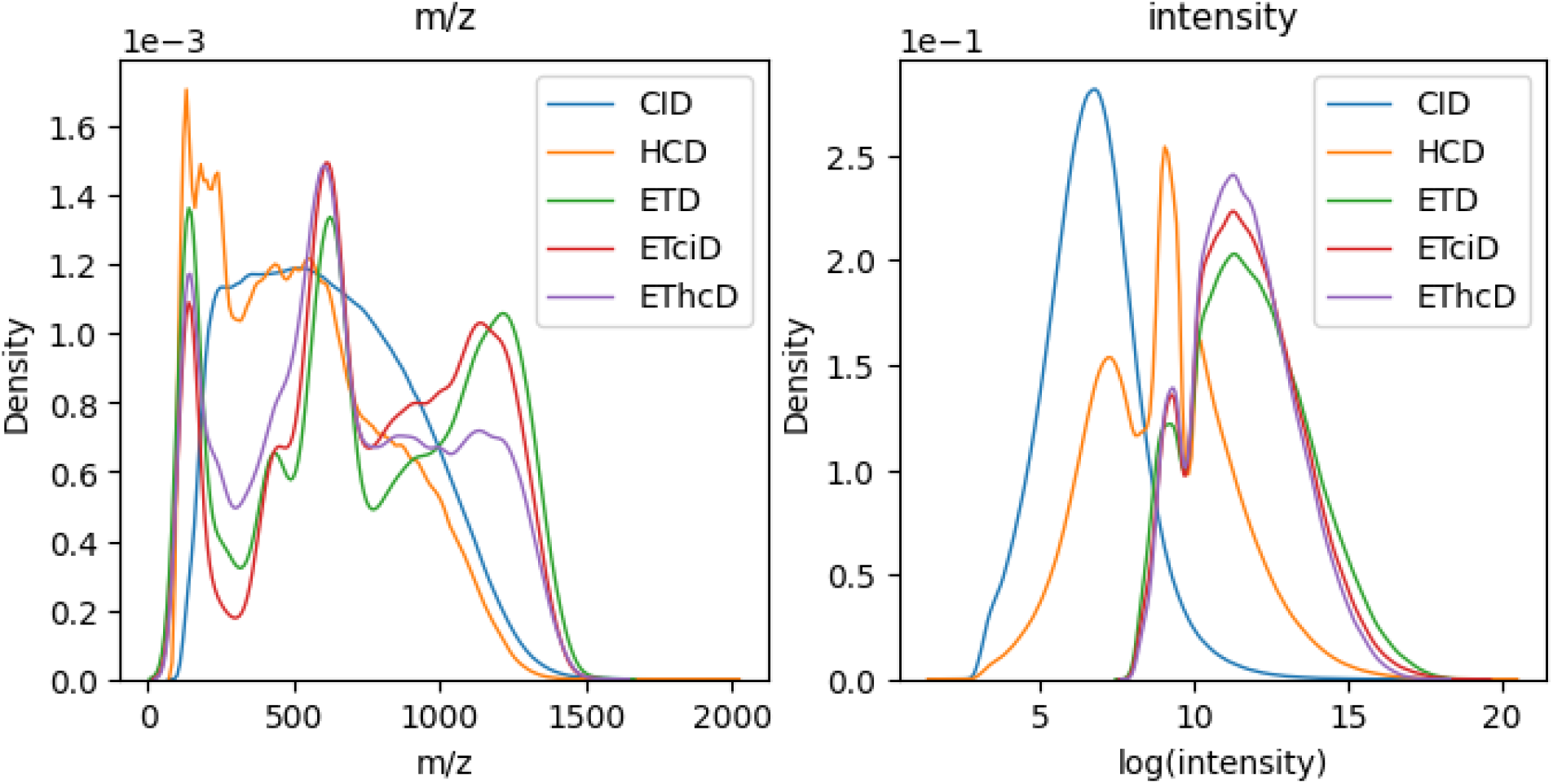
MS2 peak distribution of identical samples analyzed by different fragmentation techniques. Single sample density plot of the m/z (left) and intensity (right) peak distributions for CID, HCD, ETD, ETciD, and EThcD. Sample 02079a_BE2-TUM_isoform_50_01_01

While the three sample files from ProteomeTools display significant differences between the files, our results show that individual samples share significant similarities between fragmentation methods (Figure 10, Supplementary Figures 10-11), where we observe all ETD variants exhibiting similar distributions for both m/z and intensity values. HCD exhibits some of the same peaks and valleys as the ETD variants but is still distinct, while the CID has a completely different pattern for all files.

Along with the fragmentation technique, we also find that activation energy has a significant impact on the m/z peak distributions (Supplementary Figures 12-14). Moreover, we again observe a high correlation between ETD variants at the same reaction times, while exhibiting a significantly lower correlation within either of the variants at different activation energies. While activation energy does change the distribution of intensities, these are less impacted than m/z values.

These results show that even though two samples contain identical peptides, the fragmentation technique as well as the activation energy has a significant impact on the distribution of the reported peptide patterns. We also observe that certain methods at higher activation energies contain information about single AAs, and while these residues provide limited information for the identification of a peptide, they could be of crucial value when considering internal fragment ions in the database search. Single AA peaks and their frequencies can also contain valuable information about peptide composition and are potentially valuable for machine learning predictions. Previous attempts at using AE (specifically NCE) for ML prediction found that AE was inconsistent across files [21], that transferring between fragmentation techniques or AE limited model transferability, and that AE correction improved model performance [12].

## Conclusions

Mass spectrometry remains a powerful tool to quantify thousands of protein abundances in biological samples. Analysis of the raw experimental data is increasingly dependent on suitable computational methods [22] with a major focus on algorithms for peptide identifications and protein quantifications. However, despite a variety of different statistical, conceptual, and graph approaches, methods such as database search engines still suffer from limitations both in accuracy and run time. Novel machine learning methods hold the promise of advancing the analysis of upcoming data, as well as having a high potential for re-purposing the ample body of public data for the retrieval of valuable new biological information.

In this manuscript, we have investigated and highlighted some of the main sources of variance found within the high-throughput MS data. We identified a range of factors that increase variance in the data-generating process and analyzed the homogeneity of the variance within a project when comparing different projects. Our main finding from the statistical analyses was that global variance, which is found between projects, is significantly larger than internal variance, which is found between files in the same projects. This is exhibited through instrument settings, sample preparation, and experimental choices, all of which are significantly more homogenous within any given project, compared between projects. Furthermore, we also wanted to see how these sources of variance impacted ML capabilities by training Prosit retention time predictors on each source individually, whenever applicable.

We trained nine Prosit models, tested these models on twenty-seven datasets, and performed transfer learning fourteen times. Our findings show that training models on data most closely resembling real-life test cases is crucial, as the models’ ability to generalize outside the training data confinements are severely limited. This is illustrated by the *PT* models outperforming any other model during training and validation while having considerable performance drops when tested on randomly acquired data, or peptides not in the original m/z range.

Our results also found evidence that transfer learning can occasionally improve the performance of a pre-trained model. However, the most common scenario we observed was that models ended up mimicking non-transferred models for the same dataset, while not reducing the average amount of epochs needed for convergence. This tendency resulted in model regression in 5 of the 14 cases, and only resulted in model improvement in 1 of 14 cases. While our findings do indicate that models need to be trained on datasets from representative sources, they do not indicate that transfer learning outperforms training a new model in accuracy or computational needs (Supplementary Tables 4,5).

We argue, since a representative dataset is needed, that a research environment either has to train specialized models to their individual data collection methods or generate an unbiased dataset from publicly available data sources that attempts to mimic the intended post-training application through software such as MS2AI [19].

We further found that fragmentation spectra are rich in yet neglected information. Given the rather abundant single residue fragment ions, particularly present at higher activation energies, considerable amounts of internal ions should be present. This information has been so far mostly untapped due to the complexity to include internal ions in database search and spectrum prediction. Advanced machine learning methods might be capable of making sense of these ions despite their noisy and ubiquitous nature.

We note that a prevailing issue with the current data repositories is the missing or mislabeling of metadata. With the ongoing standardization efforts in large repositories such as PRIDE [23], this issue should fade over time. Through the analysis, we also identified the need to report more details about the experimental design, the data acquisition, and the post-processing in a comprehensive and standardized way to make them amenable as additional input for machine learning applications and thus allow for the direct training of the confounding factors.

## Methods

We use different methods to evaluate the variance caused by different setup parameters of the LC-MS experiments and their effect on ML transferability to unseen data. To assess the impact of biases and experimental heterogeneity, we trained identical deep-learning models over a range of data properties and compared their results. We used the Prosit retention time model with peptide sequence and retention time as input and output, respectively. The Prosit model architecture consists of a sequence embedding layer, a bidirectional GRU layer, and an attention layer, followed by fully connected dense layers. The retention time of all peptides in a raw file has been linearly normalized to an effective gradient; spanning between the first (0) and last (1) identified peptide to mimic the iRT calculations performed in the original paper. No further data refinement or re-annotation has been applied to the files. Initially, we followed the hyperparameter setup described in the Prosit paper but found that 32 epochs were not sufficient for model training convergence. As a result, we increased the training to 100 epochs and applied a 20-epoch patience for early stopping instead. All other parameters were identical to those described in the original paper [12]. The Prosit deep learning architecture was implemented by using the DLOmix framework [24], and all modified peptides were removed due to DLOmix constraints.

For all trained networks we sample 10% of each dataset as a hold-out test set on which all testing is conducted. The remaining data is split into training and validation sets with a ratio of 80:20. For datasets composed of multiple PRIDE projects, the hold-out datasets consist of separate projects that were randomly sampled, whereas for datasets consisting of a single project, the hold-out dataset consists of randomly sampled raw files. This provides the most accurate heterogeneous test scenarios without overlap across projects or MS runs. Furthermore, the training and validation datasets are split by peptide sequences, meaning that no peptide will be present in both the training or the validation dataset. However, since the testing data is randomly split at the project or file level, these may contain sequences that are also present in the training datasets.

The data acquisition along with filtering and model training and testing was managed using MS2AI with MongoDB in Python 3.8 with an NVIDIA v100 GPU. The data were acquired in November 2021 with the extractor API with the options “-p -mo -t 128” which allows for en-mass data acquisition from PRIDE (-p) while only fetching MaxQuant information (-mo), and increasing thread counts to 128 to allow for faster runtime (-t 128). This requires the current version of the PRIDE metadata which is downloaded using the scraper API and the -db option. The filtering was performed using the filter API using the -q option with MongoDB or string filters available in the GitHub repository. The model training and testing were performed using the network API with “-t prosit -e 100 -sos -s *n*” to train a Prosit model for 100 epochs with training and validation being split based on unique sequences and a given seed for consistent training, validation, and test splitting. When performing transfer learning, the only difference is “-t prosit-ID” which uses the weights of the trained model with the same ID. Model training times varied from 4 to 10 hours based on the size of the dataset and epochs needed to converge. All code is available in the GitHub repository along with the seeds for the runs.

We utilized a Bayesian approximation of the model uncertainty, by performing model inference with dropout enabled [25,26]. The real retention time values are then plotted against the mean predicted values, with the color of the data point corresponding to the normalized variances of the predicted values. The dropout for inference testing was applied to all layers where dropout was originally applied, with original dropout rates. This allows us to evaluate the models not just on their metric performances, but also determine the retention time ranges where the models are least certain of their predictions. This method is available in MS2AI network API using the “-id *n*” option to run *n* dropout tests and automatically generate the data visualization plots.

### 1 Single vs multi-project variance

In order to measure the difference in variance not caused by individual factors, but instead caused by systemic changes in experimental protocol, we compared the model performance of two single-project models to the performance of two multi-project models. We did this by training four Prosit models on data from four different sources: 2017 [27] and 2019 [28] ProteomeTools datasets (*PT17* and *PT19*, respectively) and two sets of acquired data from randomly sampled PRIDE projects; one limited to the 750.000 peptides filtered at 100 Andromeda score threshold, which is the score reported by MaxQuant (*Limit*), and one with 2.000.000 peptides filtered at 150 Andromeda score threshold (*Wide*). Alongside the initial training and testing, we also performed transfer learning on all models for all non-source datasets to compare their initial performance to post-transfer learning performance.

### 2 Spectra and gradient lengths

To compare and analyze the variance in gradient lengths, we extracted the metadata from each raw file (found in the *files*.*bson*.*gz*) and we extracted the gradient lengths of all runs individually, which we plotted in a histogram against the probability of each gradient length. We then calculated the cumulative distribution function of the gradient lengths for all files which we overlaid on the histogram. Then we calculated the number of unique gradients across all files from the same PRIDE accession, allowing us to visualize the variance found within projects when plotting the number of unique gradients in a histogram against the probability of each number of unique gradients.

Along with gradient length visualizations, we also trained two Prosit models to evaluate the effect of gradient length on model performance. The models were trained datasets that were divided into two groups based on their gradient lengths: short (≤ 60 minute gradients) and long (> 100 minutes gradients). The data were randomly sampled from the full 398M peptide dataset and only peptides with ≥ 150 Andromeda scores were kept. We also performed transfer learning of both gradient models to the opposing datasets. Furthermore, to test model uncertainty across the gradient, we performed model inference with dropout enabled on all four models, as described in method section 1.

### 3 m/z range filter

To visualize and compare m/z filters across files and projects, we extracted the m/z filter bounds from each raw file, and plotted the lower bounds, upper bounds and the difference between the upper and lower bounds to get the lengths. We then visualized these values in a violin plot, in order to see possible patterns or key values in the distributions.

In order to evaluate the impact of m/z filters on model performance we created two subsets of data, this time much smaller due to low peptide count; one in which all peptides lie below 360 m/z (*Lower*), and one in which all peptides lie above >1300 m/z (*Upper)*. These bounds were chosen as we are going to re-use the *PT17* and *PT19* models, which have m/z filter bounds of 360-1300 m/z, and using these datasets will allow us to evaluate peptides that are outside original filter bounds. As we did not train new models for this section, no transfer learning was applied. Again we also perform model inference with dropout enabled, in order to assess the model uncertainty for these out-of-bounds (OOB) peptides.

### 4 Fragmentation Pattern

To analyze the distribution of MS2 peaks we extracted every peak from 86 raw files [29], where we compared the raw spectra with all peaks preserved, to spectra filtered by top *n* peaks based on intensity for three values of *n*: 50, 100, and 200. MS2 spectra are often annotated or binned in order to make it fit into typical ML architectures. To illustrate how this type of binning affects the outcome distribution, we also binned each of the combined peak selected spectra at 50, 100, 200, and 500 total bins from 0-2000 m/z. We then calculated the collision ions *a, b*, and *y* as well as the ETD ions *c*, and *z* for all amino acids separately. This was done by adding -27, 1, 19, 18, and 2 mass to their single charged molecular residue weights for *a, b, y, c*, and *z* ions respectively [30,31].

Furthermore, we also sampled three representative samples from the ProteomeTools 2019 dataset: two files taken from the isoform set of tryptic peptides (*02079a_BE2-TUM_isoform_50_01_01* and *02079a_BF4-TUM_isoform_64_01_01*), and one taken from the missing protein datasets (*01974c_BH1-TUM_missing_first_8_01_01*) [32]. Each identical sample has been tested 4 times, with different fragmentation techniques and varying normalized collision energies (NCE); collision-induced dissociation (CID), high-energy C-trap dissociation (HCD), electron-transfer dissociation (ETD), electron-transfer/higher-energy collisional dissociation (EThcD) and electron-transfer/collision-induced dissociation (ETciD).

## Abbreviations

ML: Machine Learning
DL: Deep Learning
MS: Mass Spectrometry
LC-MS or MS1: Liquid Chromatography-Mass Spectrometry
LC-MS/MS, MS/MS or MS2: Tandem mass-spectrometry
m/z: Mass to charge ratio
NCE: Normalized Collision Energy
PTM: Post-translational modification
CID: Collision induced dissociation
HCD: high-energy C-trap dissociation
ETD: electron-transfer dissociation
ETciD: electron-transfer and collision-induced dissociation
EThcD: electron-transfer and higher-energy collision dissociation
PX: ProteomeXchange

## Authors’ contributions

T.G.R., K.K., R.R, V.S. conceptualized the study. T.G.R. designed and carried out the analyses with the help of K.K. T.G.R. wrote the first draft of the manuscript, with all authors contributing to the revisions. All authors read and approved the final manuscript.

## Availability of source code and requirements

Project name: MS Review Paper

Project home page: https://gitlab.com/tjobbertjob/ms-review-paper

Operating system(s): Platform independent

Programming language: Python

Other requirements: Python 3.8, MongoDB

License: e.g. GNU AGPLv3

## Data Availability

The database files, reference texts, and trained models supporting the results of this article are available in the FigShare repository [33].

