## Supplementary Figures and Tables for "Variance Analysis of LC-MS Experimental Factors and Their Impact on Machine Learning"

**Title**: Making large-scale proteomics data fit for machine learning - a data-centric perspective.

### *Table of Content*

| | |
|---|---|
| **Supplementary Figure 1.** | **1** |
| **Supplementary Figure 2.** | **1** |
| **Supplementary Figure 3.** | **2** |
| **Supplementary Figure 4.** | **2** |
| **Supplementary Figure 5.** | **3** |
| **Supplementary Figure 6.** | **3** |
| **Supplementary Figure 7.** | **4** |
| **Supplementary Figure 8.** | **5** |
| **Supplementary Figure 9.** | **5** |
| **Supplementary Figure 10.** | **6** |
| **Supplementary Figure 11.** | **7** |
| **Supplementary Figure 12.** | **7** |
| **Supplementary Figure 13.** | **8** |
| **Supplementary Figure 14.** | **9** |
| **Supplementary Figure 15.** | **10** |
| **Supplementary Figure 16.** | **11** |
| **Supplementary Table 1.** | **14** |
| **Supplementary Table 2.** | **14** |
| **Supplementary Table 3.** | **15** |
| **Supplementary Table 4.** | **15** |
| **Supplementary Table 5.** | **15** |

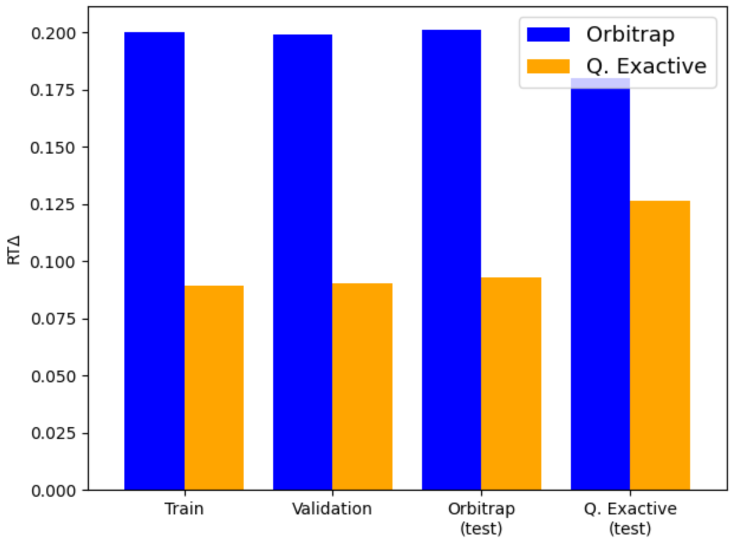


**Supplementary Figure 1.**

**Instrument model performance comparison.** Each model was trained and validated on individual internal datasets, and cross-tested on all external datasets as a basis for model comparisons. The x-axis denotes datasets, and the bars denote model performance on the given dataset.

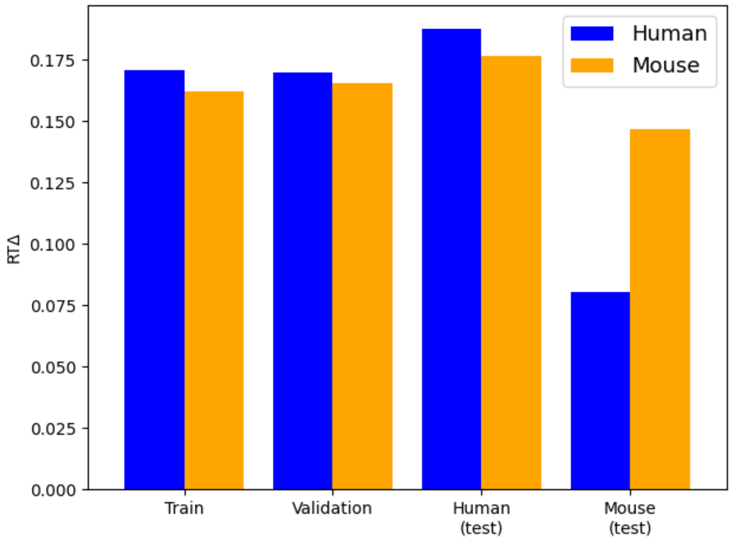


**Supplementary Figure 2.**

**Species model performance comparison.** Each model was trained and validated on individual internal datasets, and cross-tested on all external datasets as a basis for model comparisons. The x-axis denotes datasets, and the bars denote model performance on the given dataset.

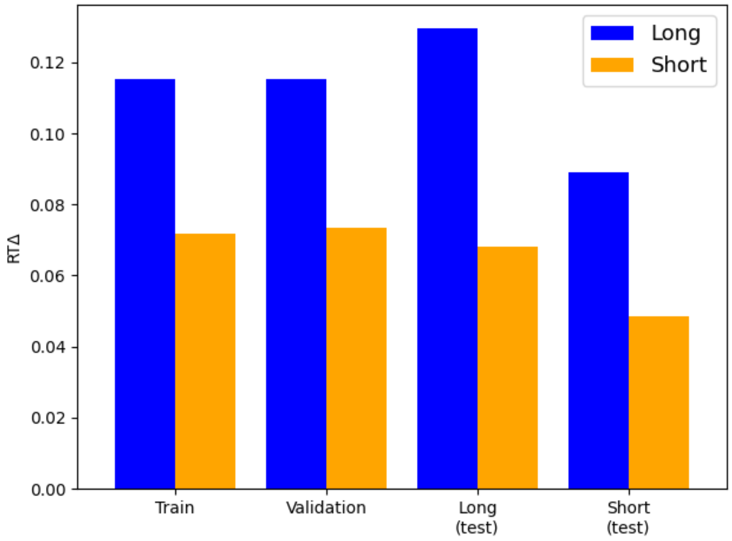


**Supplementary Figure 3.**
**Gradient model performance comparison.** Each model was trained and validated on individual internal datasets, and cross-tested on all external datasets as a basis for model comparisons. The x-axis denotes datasets, and the bars denote model performance on the given dataset.

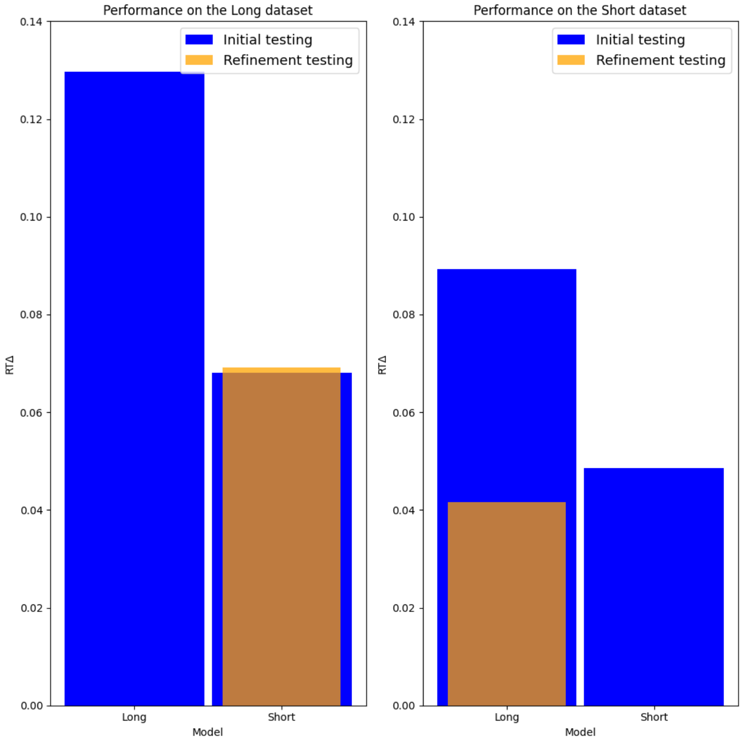


**Supplementary Figure 4.**
**Gradient based refinement model performance comparison**. Each model was refined on all previous external datasets, and re-tested on both datasets. Datasets are separated by plots, denoting the performance difference of each model when trained on, or refined to, identical datasets. Each bar has the original test metric in blue, and the refinement test metric overlaid in orange.

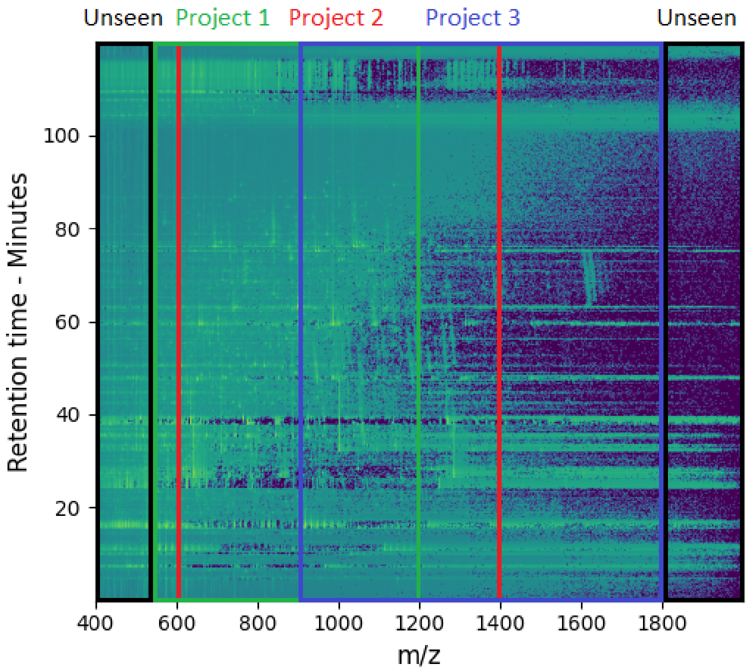


**Supplementary Figure 5.**
**Illustration of the unseen data space caused by limited search space filters.** Illustrated are three theoretical search spaces superimposed on top of a theoretical MS1 chromatogram, with unreported peptides at both ends of the data space.

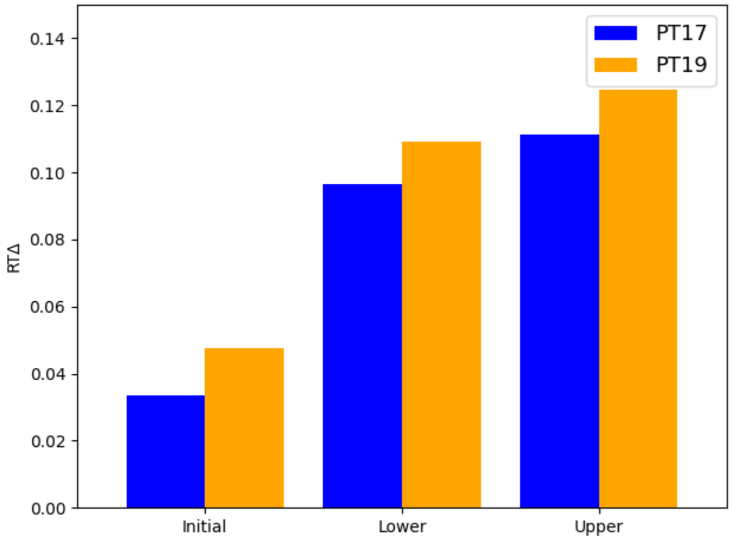


**Supplementary Figure 6.**
**Out-of-bounds performance comparison of PT models.** The *PT17* and *PT19* model comparison for their individual internal datasets (Initial), peptides with m/z below 360 (Lower) and above 1300 (Upper).

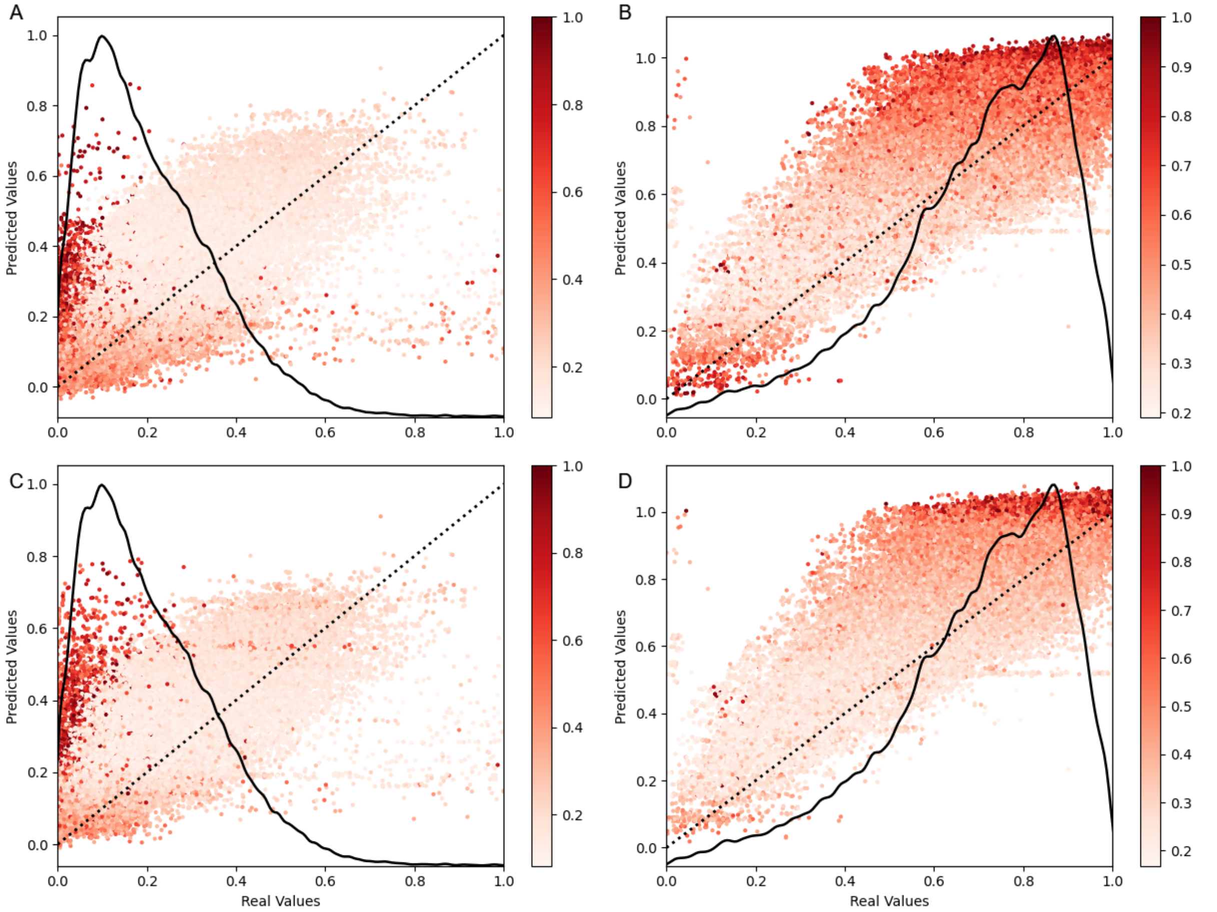


**Supplementary Figure 7.**
**Bayesian model uncertainty estimates for out-of-bounds peptides split into two datasets.** The out-of-bounds datasets are split into two datasets: peptides with m/z below 360 (Left) and above 1300 (Right). Models are separated in rows with *PT17* in the first row (A, B) and *PT19* in the last row (C, D).

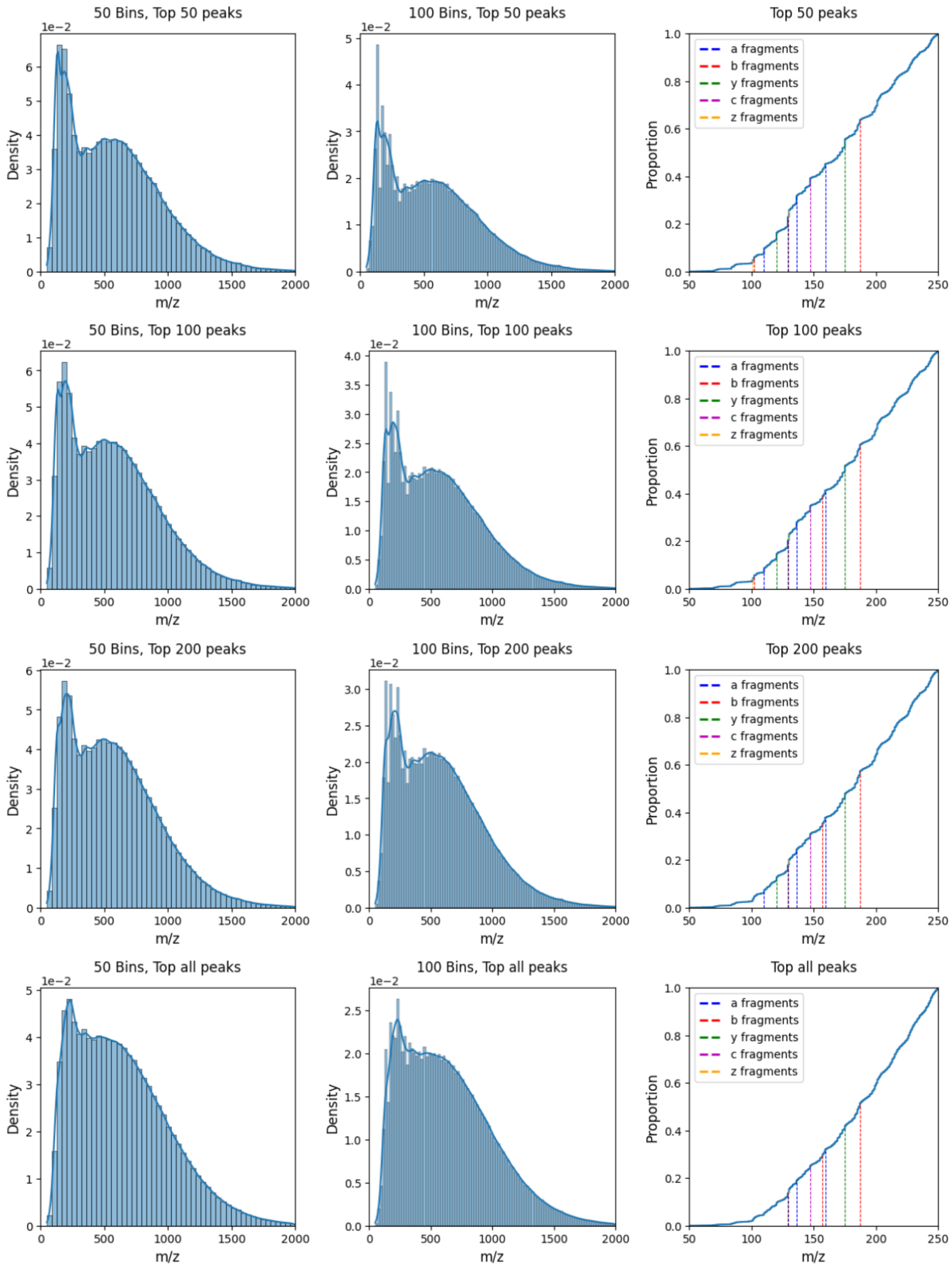


**Supplementary Figure 8.**

**Density distribution of MS2 peaks with different levels of peak picking imposed at 50 and 100 bins.** For each value of peak picking we also illustrated the cumulative distribution plot of the peaks in 50-250 m/z with single AA fragmentations overlaid

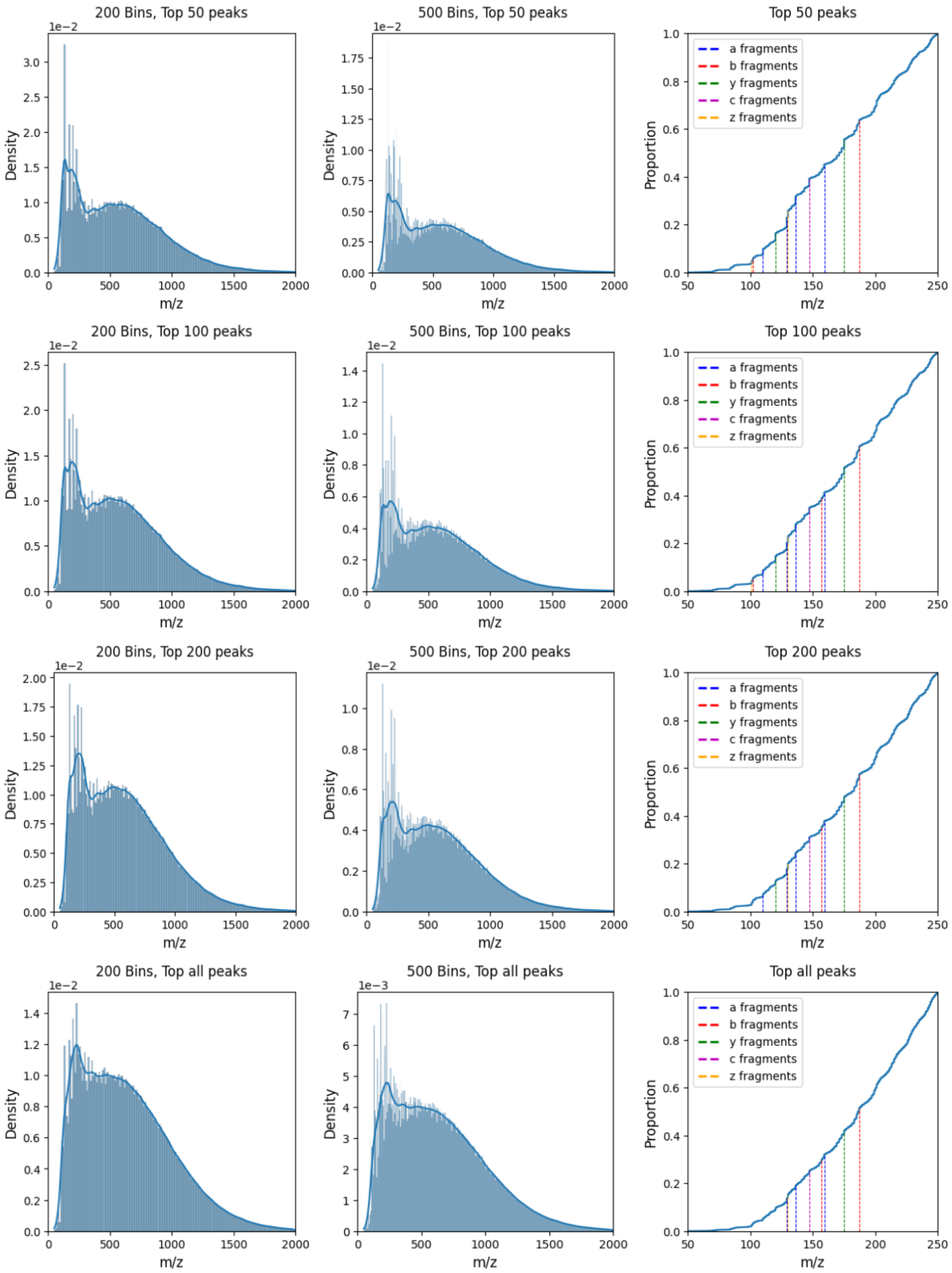


**Supplementary Figure 9.**

**Density distribution of MS2 peaks with different levels of peak picking imposed at 200 and 500 bins.** For each value of peak picking we also illustrated the cumulative distribution plot of the peaks in 50-250 m/z with single AA fragmentations overlaid

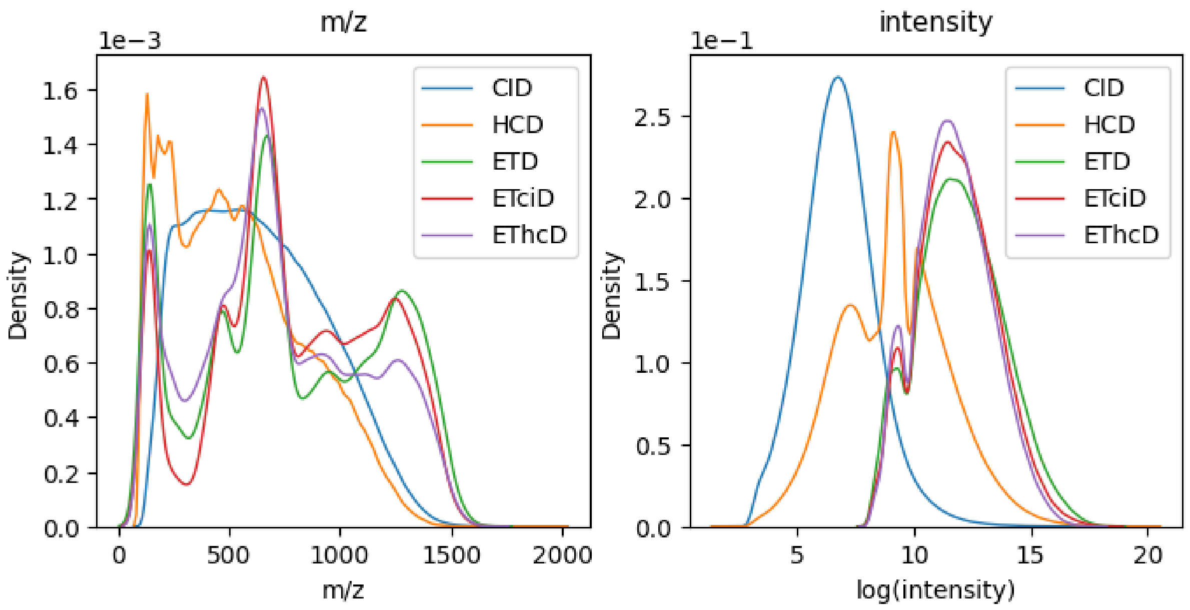


**Supplementary Figure 10.**
**MS2 peak distribution of identical samples analyzed by different fragmentation techniques.** Single sample density plot of m/z (left) and intensity (right) values of peaks for CID, HCD, ETD, ETciD, and EThcD. Sample is 02079a_BF4-TUM_isoform_64_01_01

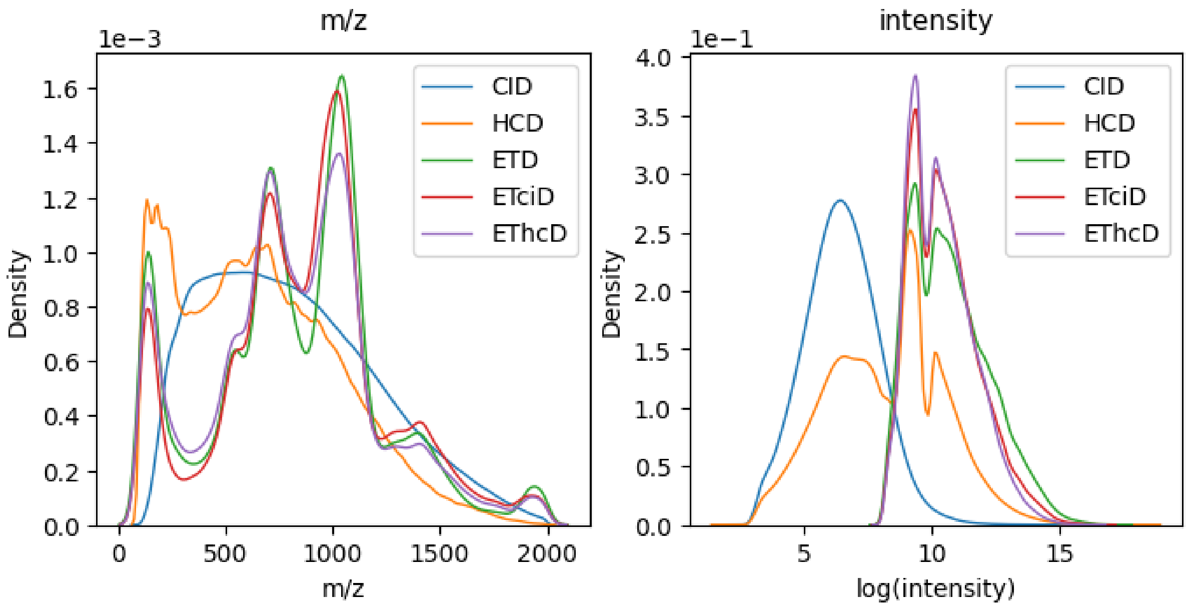


**Supplementary Figure 11.**
**MS2 peak distribution of identical samples analyzed by different fragmentation techniques.** Single sample density plot of m/z (left) and intensity (right) values of peaks for CID, HCD, ETD, ETciD, and EThcD. Sample is 01974c_BH1-TUM_missing_first_8_01_01

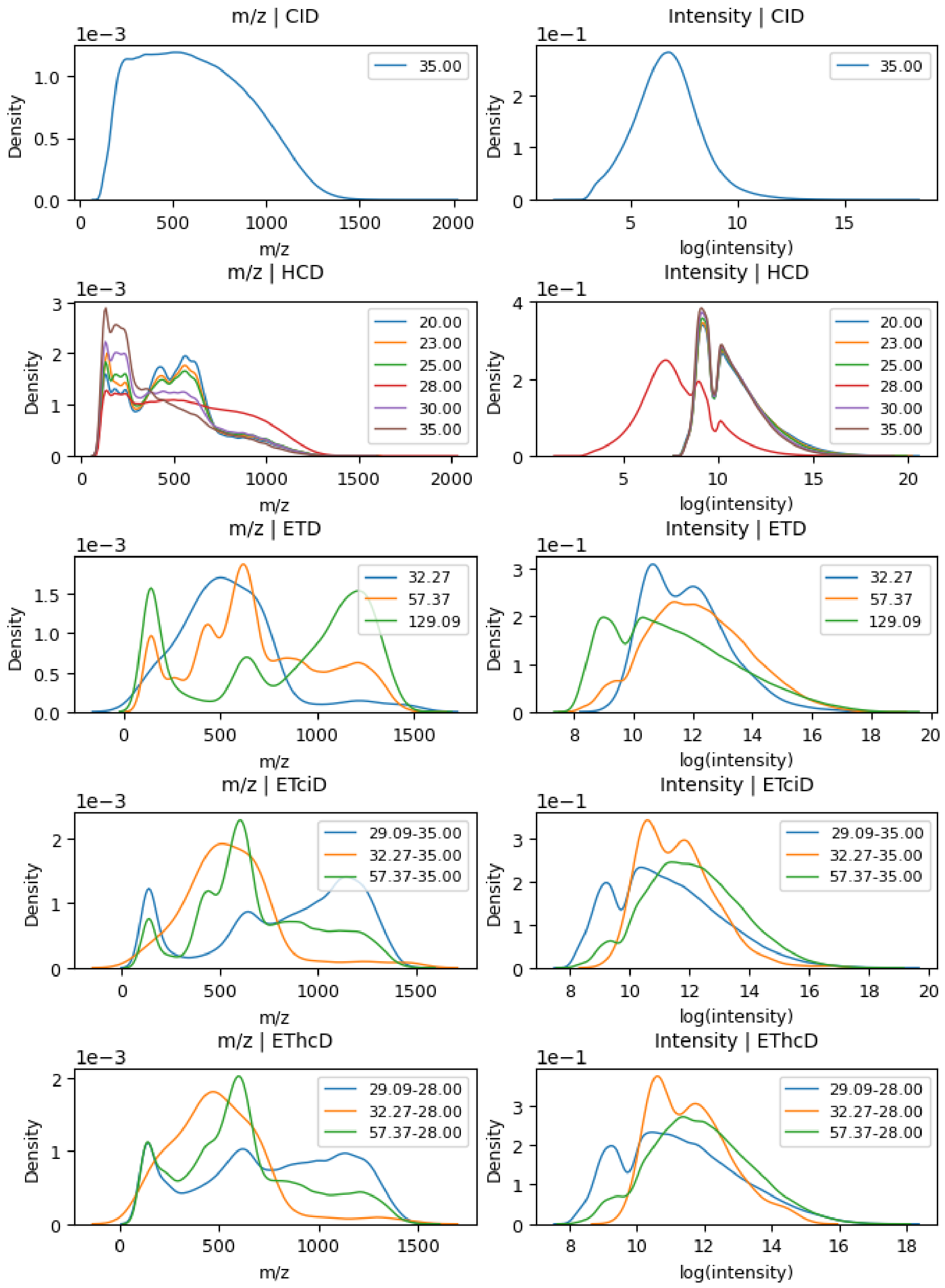


**Supplementary Figure 12.**

**Comparison of density distribution plots for peak m/z and intensity across multiple values of NCEs**. Shown is the density plot for peak m/z and intensity, stratified by the fragmentation method, and each line represents a unique NCE. This illustrates how the CE affects the peak density plots compared between fragmentation methods. Sample is 02079a_BE2-TUM_isoform_50_01_01

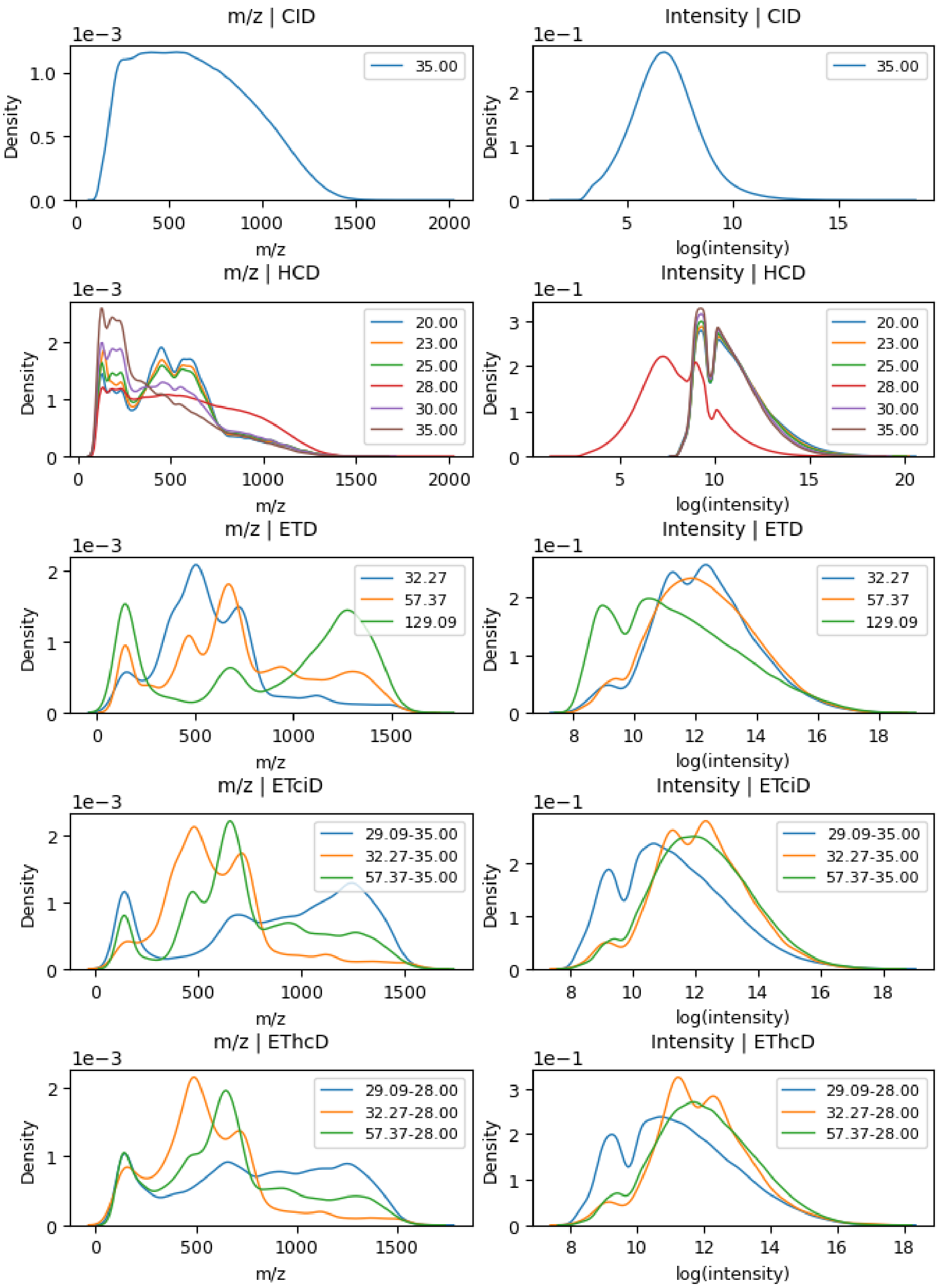


**Supplementary Figure 13.**
**Comparison of density distribution plots for peak m/z and intensity across multiple NCEs.** Shown is the density plot for peak m/z and intensity, stratified by the fragmentation method, and each line represents a unique NCE. This illustrates how the CE affects the peak density plots compared between fragmentation methods. Sample: 02079a_BF4-TUM_isoform_64_01_01

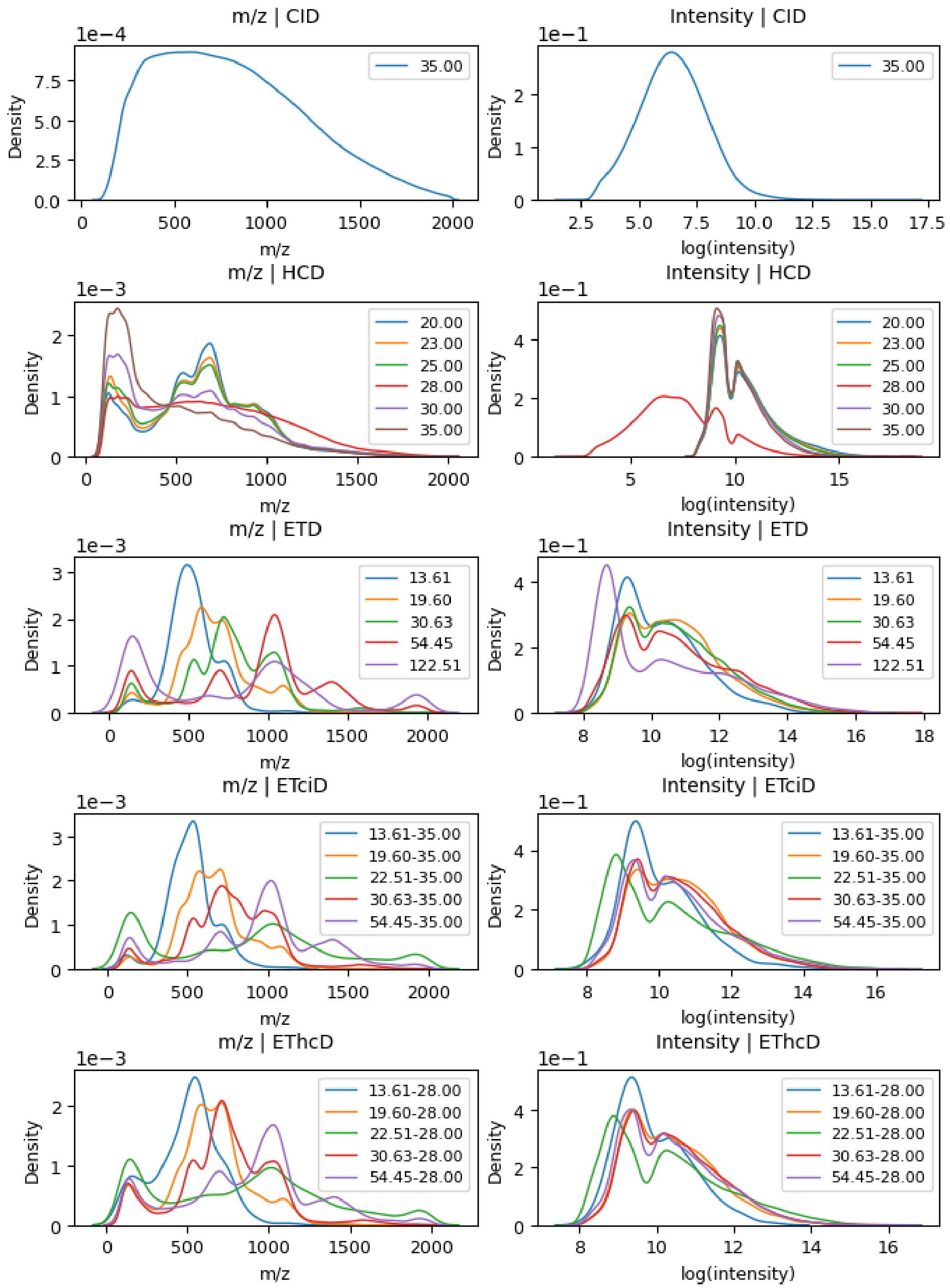


**Supplementary Figure 14.**

**Comparison of density distribution plots for peak m/z and intensity across multiple values of NCEs**. Shown is the density plot for peak m/z and intensity, stratified by the fragmentation method, and each line represents a unique NCE. This illustrates how the CE affects the peak density plots compared between fragmentation methods. Sample is 01974c_BH1-TUM_missing_first_8_01_01

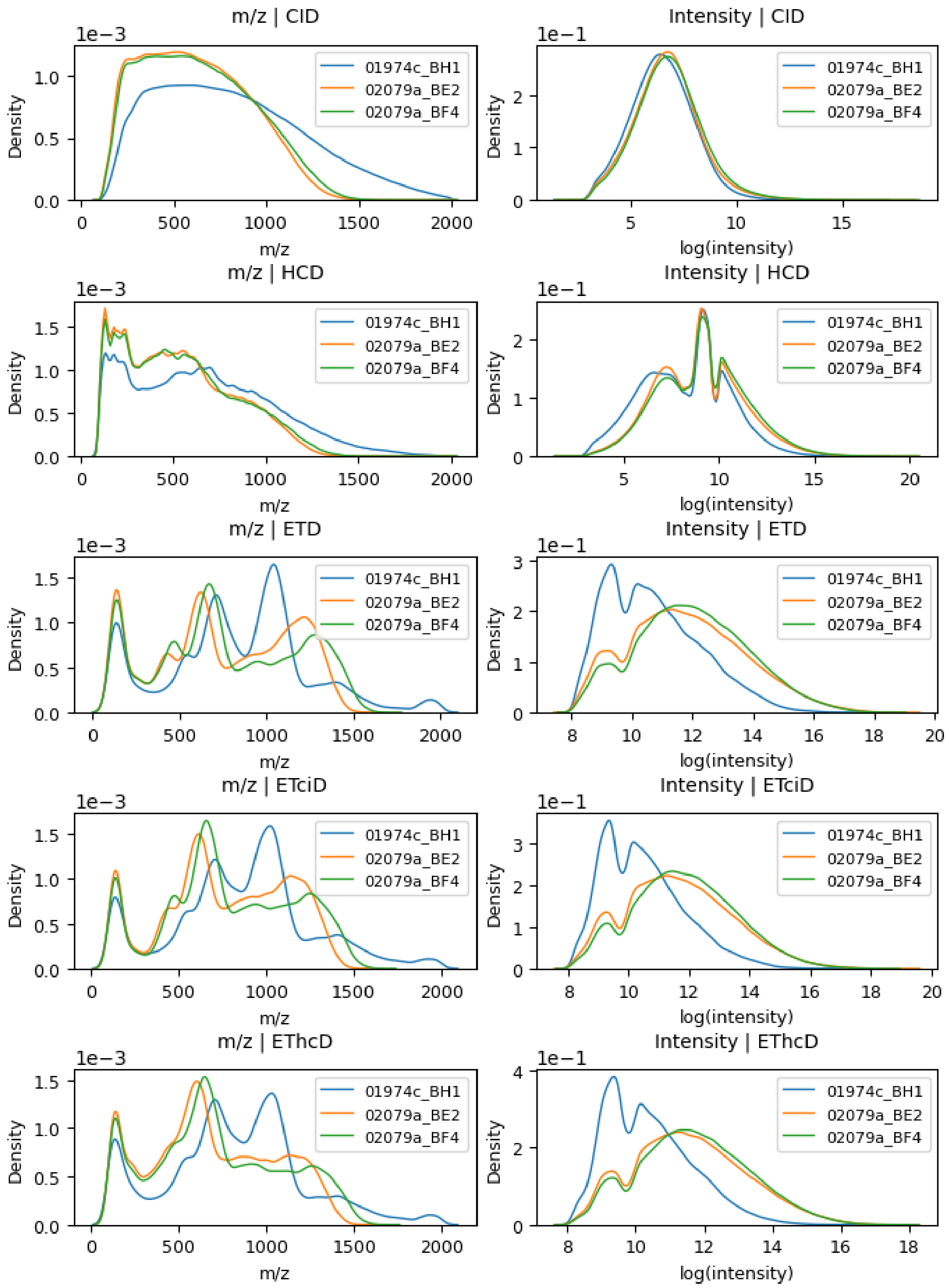


**Supplementary Figure 15.**

**Comparison of density distribution plots across files**. Each line represents one of the three files, all NCEs combined, with each row representing a separate fragmentation technique.

| m/z | Amino Acid(s) and Fragmentation | Frequencies *(50, 100, 200, all)* |
|---|---|---|
| 101.1 | Glutamine *(a)* & Lysine *(a)* | 1.15%, 1.17%, *, * |
| 102.1 | Glutamate *(a)* & Threonine *(b)* | 1.23%, 1.14%, *, * |
| 110.1 | Histidine *(a)* | 2.55%, 1.83%, 1.33%, * |
| 120.1 | Phenylalanine *(a)* & Threonine *(y)* | 3.14%, 2.17%, 1.56%, * |
| 129.1/129.2 | Glutamine *(a)* & Lysine *(b)* & Arginine *(a)* | 4.13%, 3.08%, 2.39%, 1.57% |
| 130.1 | Glutamate *(b)* & Glutamine *(z)* & Lysine *(z)* | 2.89%, 2.62%, 2.3%, 1.69% |
| 136.1 | Tyrosine *(a)* | 3.37%, 2.45%, 1.81%, 1.18% |
| 147.1 | Glutamine *(y)* & Lysine *(y)* & Glutamate *(c)* | 3.19%, 2.31%, 1.74%, 1.21% |
| 157.2 | Arginine *(b)* | *, 1.18%, 1.31%, 1.16% |
| 159.2 | Tryptophan *(a)* | 1.46%,'1.49%, 1.47%, 1.19% |
| 175.2 | Arginine *(y)* | 3.54%, '2.7%, 2.09%, '1.47% |
| 187.2 | Tryptophan *(b)* | 1.62%, 1.75%, 1.8%, 1.57% |

**Supplementary Table 1.**
**Corresponding single amino acid peak table. m/z, amino acid and fragmentation type of MS2 peaks on Figure 9.** '*' does not meet the threshold of 1% in the given *n,* and is not annotated on the CDF. *Note that 129.1 and 129.2 have been merged to one peak, due to rounding error.*

| | | Models | | | | |
|---|---|---|---|---|---|---|
| | | *PT17* | *PT19* | *Limit* | *Wide* | **Means** |
| **Datasets** | *PT17* | 100.0% | 60.7% | 119.7% | 154.1% | 108.6% |
| | *PT19* | 64.2% | 100.0% | 188.7% | 150.9% | 126.0% |
| | *Limit* | 60.7% | 77.0% | 100.0% | 41.0% | 69.7% |
| | *Wide* | 51.2% | 68.6% | 74.4% | 100.0% | 73.6% |
| | | | | | | ***94.5%*** |

**Supplementary Table 2.**
**Model epochs for training convergence overview of models trained in section r.1.**

| | | Models | | |
|---|---|---|---|---|
| | | *Long* | *Short* | **Means** |
| **Datasets** | *Long* | 100.0% | 106.9% | 103.5% |
| | *Short* | 192.9% | 100.0% | 146.5% |
| | | | | ***125.0%*** |

**Supplementary Table 3.**
**Model epochs for training convergence overview of models trained in section r.2.**

| | | Models | | |
|---|---|---|---|---|
| | | *Human* | *Mouse* | **Means** |
| **Datasets** | *Human* | 100.0% | 103.1% | 101.6% |
| | *Mouse* | 45.3% | 100.0% | 72.7% |
| | | | | ***87.1%*** |

**Supplementary Table 4.**
**Model epochs for training convergence overview of models trained for organisms.**

| | | Models | | |
|---|---|---|---|---|
| | | *Orbitrap* | *Q. Exactive* | **Means** |
| **Datasets** | *Orbitrap* | 100.0% | 156.4% | 128.2% |
| | *Q. Exactive* | 157.1% | 100.0% | 128.6% |
| | | | | ***128.4%*** |

**Supplementary Table 5.**
**Model epochs for training convergence overview of models trained for instruments.**
